# The transcription factor Xrp1 orchestrates both reduced translation and cell competition upon defective ribosome assembly or function

**DOI:** 10.1101/2021.07.12.452023

**Authors:** Marianthi Kiparaki, Chaitali Khan, Virginia Folgado Marco, Jacky Chuen, Nicholas E. Baker

## Abstract

Ribosomal Protein (*Rp*) gene haploinsufficiency affects overall translation rate, leads to cell elimination by competition with wild type cells in mosaic tissues, and sometimes leads to accumulation of protein aggregates. The changes in ribosomal subunit levels observed are not sufficient for these effects, which all depend on the AT-hook, bZip domain protein Xrp1. In *Rp^+/−^* cells, Xrp1 reduced global translation through PERK-dependent phosphorylation of eIF2α. eIF2α phosphorylation was sufficient to reduce translation in, and also enable cell competition of, otherwise wild type cells. Unexpectedly, however, many other defects reducing ribosome biogenesis or function (depletion of TAF1B, eIF2, eIF4G, eIF6, eEF2, eEF1α1, or eIF5A), also increased eIF2α phosphorylation and enabled cell competition. In all cases this was through the Xrp1 expression that was induced, placing Xrp1 as the downstream instigator of cell competition that also contributed to overall translation deficits. In the absence of Xrp1, translation differences between cells were not themselves sufficient to trigger cell competition. Thus, Xrp1, which is shown here to be a sequence-specific transcription factor, is the master regulator that triggers cell competition and other consequences of multiple ribosomal stresses.

## INTRODUCTION

It would be difficult to exaggerate the importance of ribosomes. Eukaryotic ribosomes comprise 4 rRNAs and 80 proteins combined into Large and Small subunits (LSU and SSU) that, together with multiple initiation and elongation factors, constitute the translational apparatus for protein synthesis(Jackson, Hellen, & Pestova, 2010; Thomson, Ferreira-Cerca, & Hurt, 2013). Ribosome biogenesis and the regulation of translation are important targets of cellular regulation, and defects affecting ribosomes and translation are implicated in many diseases, from neurodegeneration to cancer(Aspesi & Ellis, 2019; Hetman & Slomnicki, 2019)(Genuth & Barna, 2018; Ingolia, Hussmann, & Weissman, 2019; Joazeiro, 2019; Phillips & Miller, 2020). Mutations affecting rRNA synthesis, ribosomal protein genes (Rp genes), and some other ribosome biogenesis factors give rise to ribosomopathies, a family of translation-related diseases(Kampen, Sulima, Vereecke, & De Keersmaecker, 2020). The ribosomopathy Diamond Blackfan Anemia (DBA) most commonly results from heterozygosity for mutations in Rp genes, and is characterized by early onset anemia, cancer predisposition, and sometimes diminished growth and skeletal defects(Draptchinskaia et al., 1999; Choesmel et al., 2007; Danilova & Gazda, 2015; Da Costa, Narla, & Mohandas, 2018). Most ribosomal protein genes are haploinsufficient in *Drosophila* also, where their dominant ‘Minute’ phenotype was named by Bridges and Morgan on account of the small, thin cuticular bristles observed, in addition to developmental delay(Bridges & Morgan, 1923; Lambertsson, 1998; Marygold et al., 2007).

Rp gene loci were recently proposed to be important indicators of aneuploidy(Ji, Chuen, Kiparaki, & Baker, 2021). It was known that aneuploid cells can be selectively eliminated from embryonic and developing mammalian tissues, but the mechanisms responsible have been uncertain(Bolton et al., 2016; McCoy, 2017). In *Drosophila*, cells heterozygous for mutations in *Rp* genes are selectively eliminated from mosaic imaginal discs, where they are replaced by neighboring wild type cells(Morata & Ripoll, 1975; Simpson, 1979). This phenomenon, named ‘cell competition’, represents a process whereby cells that present differences from their neighbors can be eliminated from growing tissues, thought to enable the removal of cells that might be deleterious to the tissue(Morata & Ripoll, 1975; Lawlor, Perez-Montero, Lima, & Rodriguez, 2019; Baker, 2020; Vishwakarma & Piddini, 2020; Marques-Reis & Moreno, 2021; Morata, 2021). Because *Rp* gene dose is likely to be affected whenever one or more chromosomes or substantial chromosome regions are monosomic, cell competition could help eliminate aneuploid cells on the basis of altered *Rp* gene dose(McNamee & Brodsky, 2009). This mechanism was recently demonstrated to occur in *Drosophila* imaginal discs(Ji et al., 2021). Such a role of cell competition is potentially significant for tumor surveillance, since tumors almost always consist of aneuploid cells, and for healthy aging, since aneuploid cells accumulate during aging(Hanahan & Weinberg, 2011; Lopez-Otin, Blasco, Partridge, Serrano, & Kroemer, 2013). In addition to their mutation in DBA, this provides another reason why it is important to understand the cellular effects of *Rp* mutations, and how they lead to cell competition.

Unsurprisingly, *Rp* mutant heterozygosity generally leads to reduced translation(Boring, Sinervo, & Schubiger, 1989; Oliver, Saunders, Tarle, & Glaser, 2004). It might be expected that a 50% reduction in ribosome subunit biogenesis would be responsible, but remarkably, in *Drosophila* this and many other features of *Rp* haploinsufficiency, including cell competition in the presence of wild type cells, depend on a bZip, AT-hook putative transcription factor encoded by the *Xrp1* gene(Lee et al., 2018). *Xrp1* is responsible for >80% of the transcriptional changes that are seen in *Rp^+/−^* wing imaginal discs(Lee et al., 2018). Thus, reduced translation, which is a feature of Rp haploinsufficiency from yeast to mice and humans, may have a transcriptional basis(Lee et al., 2018). Accordingly, we could detect only modest reductions in SSU concentration in heterozygous *RpS3*, *RpS17* or *RpS18* mutants, although *RpL27A* haploinsufficiency reduced steady state LSU numbers by ∼30% (Lee et al., 2018). Some of these findings now have support from yeast studies, where deletion of single *Rp* loci present in paralogous pairs (a recent genome duplication has left yeast with many such *Rp* gene pairs) potentially mimics heterozygosity for a single copy gene in diploid organisms. The large majority of translational changes described by ribosome profiling of such pseudo-heterozygotes turned out to reflect changes in mRNA abundance, implicating a predominantly transcriptional response to *Rp* mutations in yeast also(Cheng et al., 2019). Mass spectrometry and rRNA measurements of the yeast strains further suggested that ribosome numbers were little affected in most *RpL* gene deletion strains, whereas some *RpS* deletions increased LSU concentrations by up to 1.5x(Cheng et al., 2019).

These findings raise many mechanistic questions. How does *Rp* haploinsufficiency activate *Xrp1* gene expression? How does this putative transcription factor control overall translation, if not through altered ribosome numbers? Are differences in translation rate between cells the cause of cell competition, or is cell competition due to other consequences of Xrp1 activity?

Alternative views of the *Rp* mutant phenotype have also been presented. Aside from the idea that reduced ribosome levels alter translation directly and are predominantly responsible for human DBA(Mills & Green, 2017; Khajuria et al., 2018), two recent studies propose that degradation of excess orphan Rp suppresses proteasome and autophagic flux in *Drosophila Rp* mutants, leading to protein aggregation and proteotoxic stress. They propose that proteotoxic stress suppresses translation, and renders *Rp*^+/−^ cells subject to competition with wild type cells through a further oxidative stress response(Baumgartner, Dinan, Langton, Kucinski, & Piddini, 2021; Recasens-Alvarez et al., 2021). This view does not propose any role for the Xrp1 protein, or for transcriptional regulation of translation or cell competition. In addition, in concluding that autophagy is protective for *Rp* mutant cells (Baumgartner et al., 2021; Recasens-Alvarez et al., 2021), these studies contradict previous conclusions that autophagy is only increased in *Rp* mutant cells next to wild type cells, where it promotes cell death(Nagata, Nakamura, Sanaki, & Igaki, 2019).

Here we further investigate the basis of the *Rp* mutant phenotype in *Drosophila*. The results reaffirm the central role of Xrp1 in multiple aspects of the *Rp* mutant phenotype. We confirm the modest effects of *Rp* haploinsufficiency on numbers of mature ribosome subunits, and show directly that ribosome precursors accumulate in *Rp* mutants. We find that translation is reduced in *Rp* mutant cells through eIF2α phosphorylation, but both this and the protein aggregation observed (which appears specific for mutations affecting SSU proteins) require Xrp1 and so are not direct post-transcriptional consequences of ribosome assembly defects. We report that interfering with translation, whether through eIF2α phosphorylation or by multiple other routes disrupting ribosome assembly or function, can subject otherwise wild type cells to competition with normal cells. This is not because translation differences between cells cause cell competition directly, however, but because defects in both ribosome biogenesis and function that affect translation are all found to activate Xrp1, which then mediates the cell competition engendered by these translational stresses. Without Xrp1, translation differences are insufficient for cell competition. We then show that Xrp1 is a sequence-specific transcription factor that is required for cell competition in response to multiple triggers and is responsible for multiple aspects of the *Rp* mutant phenotype, potentially including transcription that has previously been taken as reporters of oxidative stress. Altogether, these studies clarify discrepancies in previously published work, and refocus attention on transcriptional responses to ribosome and translation defects mediated by Xrp1, with implications for the mechanisms and therapy of multiple ribosomopathies, and for the surveillance of aneuploid cells.

## RESULTS

### Ribosome levels in *Rp*^+/−^ cells

Abnormal cellular levels of ribosome subunits are proposed to affect translation in ribosomopathies(Mills & Green, 2017). Multiple models of DBA accordingly seek to reduce steady-state Rp concentration to 50% of normal(Heijnen et al., 2014; Khajuria et al., 2018). By measuring *Drosophila* rRNA levels in northern blots, however, we had previously concluded that whereas cellular levels of ribosome subunits were affected in heterozygotes for an *RpL27A* mutant, multiple *Rp* mutations affecting SSU proteins led only to ∼10% reduction in SSU levels that was not statistically significant(Lee et al., 2018). A caveat to this conclusion was the use of tubulin mRNA and actin mRNA as loading controls. While mRNA-seq shows that the proportions of actin and tubulin mRNAs are not much affected in *Rp*^+/−^ genotypes(Kucinski, Dinan, Kolahgar, & Piddini, 2017; Lee et al., 2018), it could be that total mRNA amounts are altered by *Rp* mutations, which would affect the conclusions regarding rRNA. In bacteria, it is well-established that ribosomes protect mRNA from turnover, so that reduced ribosome numbers reduce overall mRNA levels as well(Yarchuk, Jacques, Guillerez, & Dreyfus, 1992; Hui, Foley, & Belasco, 2014). The situation in eukaryotic cells may not be the same as in bacteria(Belasco, 2010). Still, we decided to measure rRNA levels again using a non-coding RNA as loading control. We chose the 7SL RNA component of Signal Recognition Particle, an abundant non-coding RNA that is expressed in all cells.

Changes in LSU and SSU levels inferred from 5.8S and 18S rRNA abundance, normalized to 7SL RNA levels, are shown in Figure 1, and a representative northern blot in Figure 1A. Similar to what was observed previously, Xrp1 mutations had no effect on apparent LSU or SSU levels in the wild type or in heterozygotes for any of four mutant loci, *RpS18*, *RpS3*, *RpL27A*, and *RpL14*, reaffirming that Xrp1 is unlikely to affect translation rate through an effect on ribosome subunit concentrations (Figure 1B,C). Accordingly, *Xrp1^+/+^* and *Xrp1^+/−^* data were combined together to compare the effects of *Rp* mutations. We confirmed that LSU numbers were reduced in the *RpL27A* mutant, and extended this observation to mutation in a second RpL gene, *RpL14* (Figure 1D). Unlike our previous study, SSU levels were reduced 20-30% in *RpS18*, *RpS3* and *RpL14* mutants when normalized to the non-coding 7SL RNA, and these reductions were significantly different from the control (Figure 1E). By contrast, *RpL27A* did not change SSU numbers (Figure 1E).

**Figure 1.**
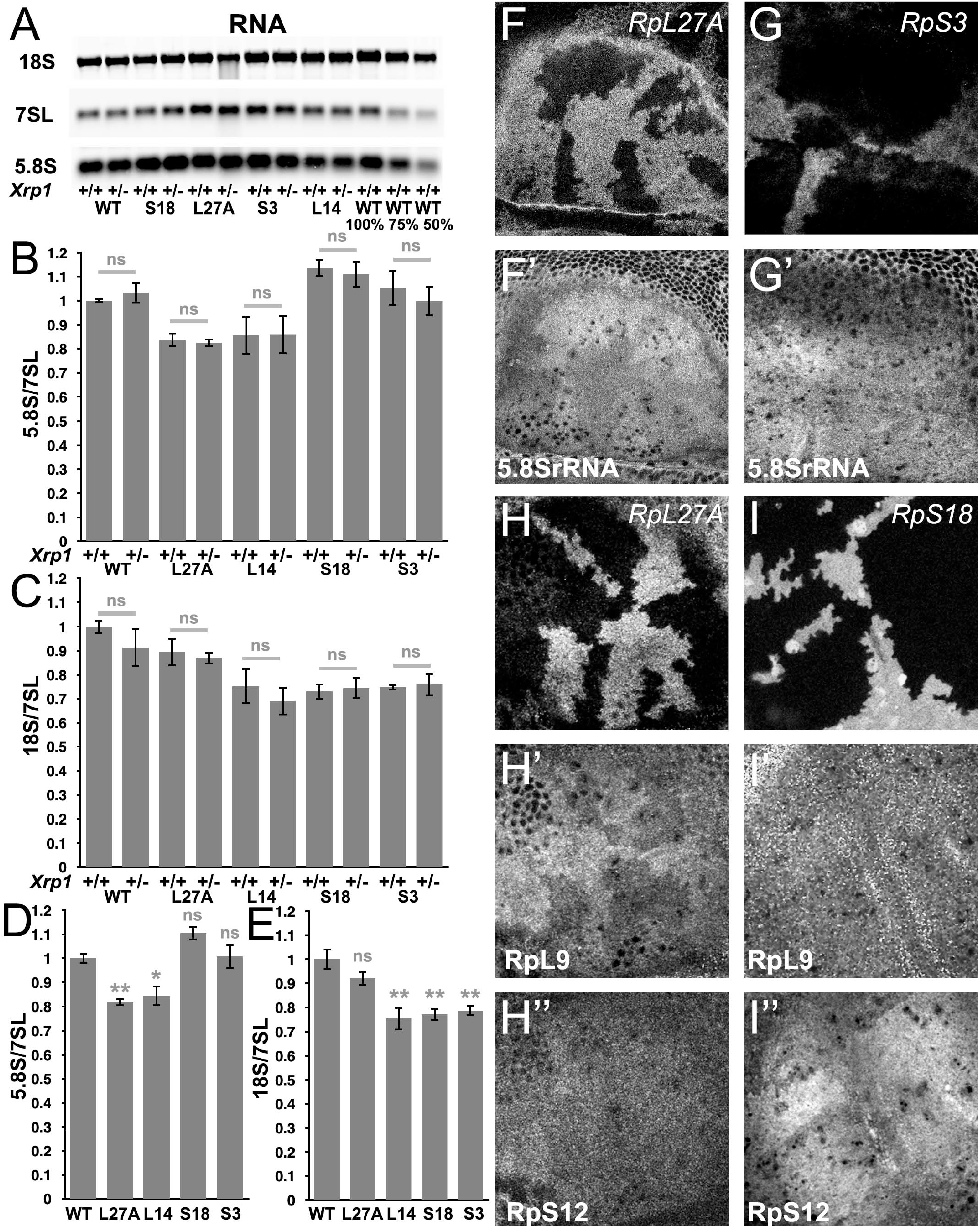
Modest changes in ribosomal subunit concentrations in *Rp* mutant wing discs. A) Similar amounts of wing disc RNA from indicated genotypes separated and transferred for northern blotting with, in this case, probes specific for the 18S rRNA of the ribosomal SSU, the 7SL non-coding RNA for the Signal Recognition Particle, and the 5.8S rRNA of the ribosomal LSU. Right-most two lanes show serial two-fold dilutions of the wild type sample. Panels B-E show signal quantification from multiple such northerns. B) *Xrp1* mutation did not affect LSU concentration in any *Rp* genotype. Significance shown only for *Xrp1^+/+^* to *Xrp1^+/−^* comparisons between otherwise similar genotypes. Exact Padj values were: 6.05, 5.16, 1.93, 6.37, 2.62 respectively. C) *Xrp1* mutation did not affect SSU concentration in any *Rp* genotype. Significance shown only for *Xrp1^+/+^* to *Xrp1^+/−^* comparisons between otherwise similar genotypes. Exact Padj values were: 4.70, 5.95, 6.94, 4.41, 8.05 respectively. D) Two *RpL* mutations reduced LSU concentrations. Significance shown only for comparisons between mutant genotypes and the wild type. Exact Padj values were: 0.00423, 0.0117, 0.0877, 0.858 respectively. E) Two *RpS* mutations, as well as *RpL14*, reduced SSU concentrations. Significance shown only for comparisons between mutant genotypes and the wild type. Exact Padj values were: 0.135, 0.000218, 0.000395, 0.000602 respectively. Panels F-I show comparisons between antibody labelings of 5.8S rRNA, anti-RpL9, or anti-RpS12 between wild type and *Rp^+/−^* cells in mosaic wing imaginal discs. F,F’) *RpL27A* mutation reduced levels of 5.8SrRNA. G,G’) *RpS3* mutation had negligible effect on 5.8S rRNA levels. H,H’,H”) *RpL27A* mutation reduced levels of the LSU component RpL9 but not of the SSU component RpS12. I,I’,I”) *RpS18* mutation reduced levels of the SSU component RpS12 but not of the LSU component RpL9. Statistics: One-way Anova with Bonferroni-Holm multiple comparison correction was performed for panels B-E, which were each based on 3 biological replicates. ns - p≥0.05. * - p<0.05. ** - p<0.01.

To confirm these findings using an independent method, we performed tissue staining with a monoclonal antibody, mAbY10B, that recognizes rRNA, and particularly a structure in the 5.8S rRNA that is part of the LSU(Lerner, Lerner, Janeway, & Steitz, 1981). Consistent with Northern analysis, immunostaining of mosaic wing imaginal discs confirmed lower 5.8S rRNA levels in *Rp27A*^+/−^ cells compared to *Rp27A*^+/+^ cells in the same wing discs (Figure 1F, Figure 1-figure supplement 1A). By contrast, no reduction in mAbY10B staining was observed in cell mutated for either of two SSU components, *RpS3* or *RpS17*, consistent with the northern blot measurements of 5.8S rRNA levels (Figure 1G, Figure 1-figure supplement 1B-D).

To gain further support for these findings, we compared Rp protein levels by immunostaining mutant and control cells in the same imaginal discs. We used antibodies against RpL9 and RpL10Ab as markers for LSU, and against RpS12 and RACK1 as markers for SSU. *RpL27A* mutant cells contained lower levels of LSU protein, and slightly lower levels of SSU protein(Figure 1H, Figure 1-figure supplement 2A).. *RpS3*, *RpS17*, and *RpS18* mutant cells contained lower levels of the SSU protein, and *RpS18* slightly higher levels of the LSU protein RpL10Ab, even in the Xrp1 mutant background (Figure 1I, Figure 1-figure supplement 2B, C-F). These tissue stainings qualitatively support the conclusion that levels of SSU components are generally reduced in *RpS*+/− cells and *RpL27A+/−* cells, whereas LSU levels are only reduced in *RpL27A+/−* cells, in comparison to wild type cells within the same preparation, and that these changes are modest and unaffected by Xrp1, even though *Xrp1* mutation restores normal global translation rate(Lee et al., 2018).

### Ribosome precursors accumulate in *Rp^+/−^* cells

An additional, or alternative, potential effect of *Rp* mutations is the accumulation of unused ribosome precursors and assembly intermediates. In yeast, depleting almost any Rp arrests ribosome biogenesis at some stage, reflecting individual requirements for ribosome assembly(Ferreira-Cerca, Poll, Gleizes, Tschochner, & Milkereit, 2005; Poll et al., 2009) (Ferreira-Cerca et al., 2005; Ferreira-Cerca et al., 2007; Woolford & Baserga, 2013; Henras, Plisson-Chastang, O’Donohue, Chakraborty, & Gleizes, 2015). *Rp* haploinsufficiency might delay biogenesis at these same steps, perhaps leading to accumulation of particular precursor states. To assess ribosome biogenesis in *Rp^+/−^* mutants, intermediates were quantified by Northern blotting using probes specific for sequences that are excised from the rRNA as the ribosome assemble and mature. In *Drosophila*, two parallel pathways A and B excise ITS1, ITS2, and the N-terminal EXT sequences, and process the resulting rRNAs, until the mature 28S (processed into 28Sa and 28Sb in *Drosophila*), 18S and 5.8S rRNAs are produced by the end of ribosome biogenesis (Figure 2A)(Long & Dawid, 1980). We used specific probes to identify rRNA intermediates on northern blots (Figure 2A-D; Figure 2 Supplement 1). As predicted, intermediates accumulated in each of the *Rp^+/−^* genotypes (see Figure 2 legend for details). These findings support the idea that *Rp* gene haploinsufficiency leads to ribosome biogenesis delays, and corresponding accumulation of assembly intermediates.

**Figure 2.**
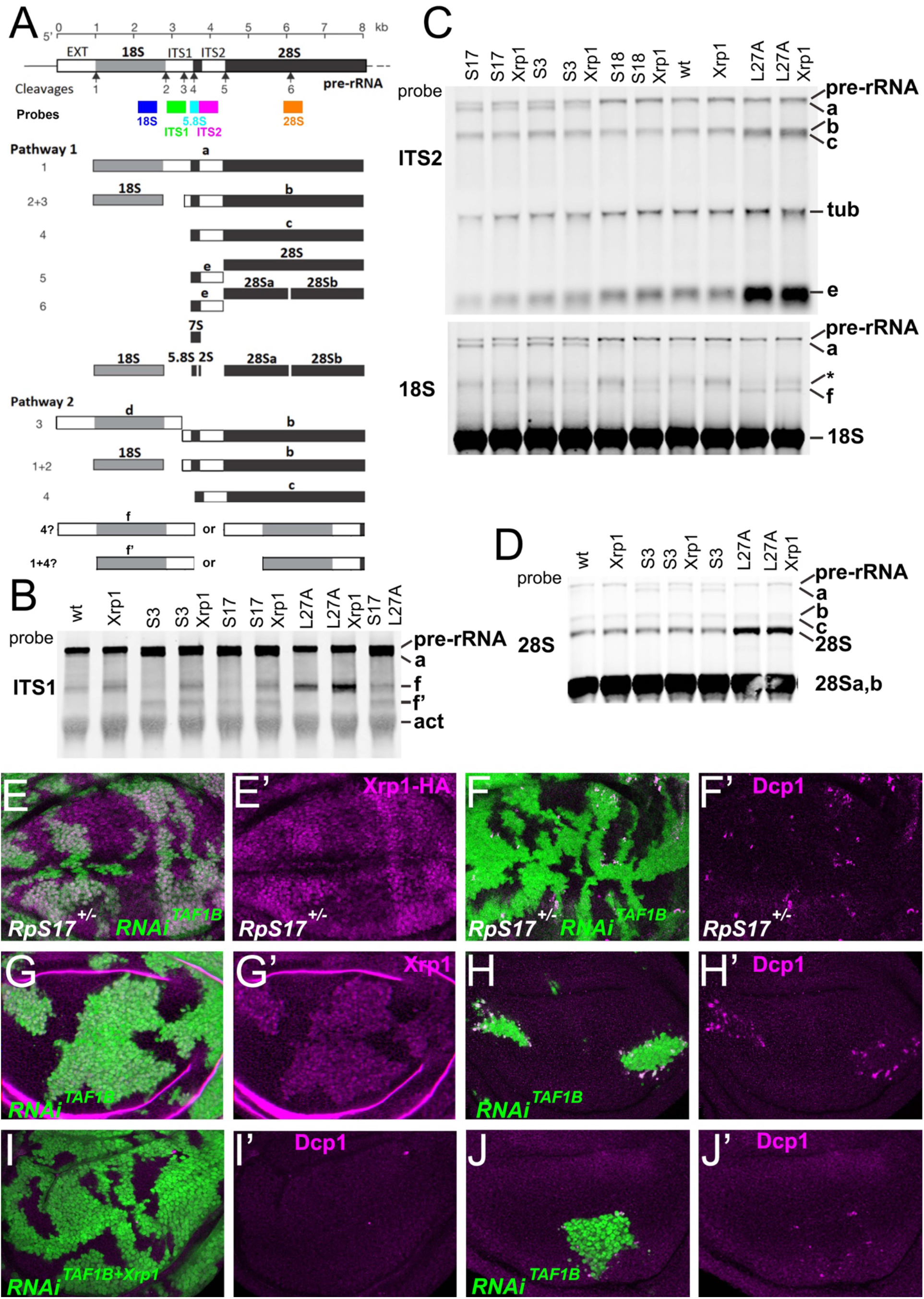
Ribosome biogenesis defects and their consequences. A) Two pathways of rRNA processing and the intermediates that result were characterized in *D.* melanogaster embryos by Long and Dawid. Mature 18S, 5.8S and 28Sa,b rRNAs are processed from the pre-RNA, along with the removal of two interval sequences ITS1 and ITS2. The cleavages sites were described by Long and Dawid. Colored boxes indicate the probes used in the present study. The 5.8S probe overlaps with 147 bases at 3’of the ITS1 region, excluding cleavage site 3. Additional intermediates **f** and **f’** were observed in the wing imaginal disc samples. These were recognized by ITS1, 5.8S (Figure 2 Supplement 1) and 18S probe and therefore extended beyond the cleavage site 3, although whether beyond site 4 was uncertain. B-D) Northern blots of total RNA purified from wild-type and Rp+/− wing discs, probed as indicated. B) Reprobed with ITS1 after an initial actin probe. C) Reprobed with ITS2 and then 18S probes after an initial tubulin probe. Intermediates b, f and the 28S rRNA (which in *Drosophila* is a precursor to the mature 28Sa and 28Sb rRNAs) were detected in wild type and *Xrp1^+/−^* wing discs, other intermediates only in *Rp^+/−^* genotypes. *RpS3^+/−^* and *RpS17^+/−^* had lower levels of pre-RNA and intermediate **f** but accumulate intermediates **a** and **f’**, which might indicate delays in cleavages 2 and 3. *RpS18^+/−^* had increased levels of pre-RNA and intermediate **f**. *RpL27A^+/−^* accumulated bands **b**, **c**, **e**, and **f** and 28S. The effect on **f** suggests crosstalk between RpL27A and SSU processing. E-I) show single confocal planes from mosaic third instar wing imaginal discs E) TAF1B depletion (green) increased Xrp1-HA levels in *RpS17*^+/−^ discs (magenta, see also E’). F) TAF1B depletion (green) increased in *RpS17*^+/−^ discs led to cell death at the boundaries with undepleted cells (active Dcp1 staining in magenta, see also F’). G) TAF1B depletion (green) also increased Xrp1 protein levels in *RpS17*^+/+^ discs (magenta, see also G’). H) TAF1B depletion (green) led to cell death at the boundaries with undepleted cells (active Dcp1 staining in magenta, see also H’). I) Co-depletion of Xrp1 with TAF1B (green) largely abolished cell death at the clone interfaces (active Dcp1 staining in magenta, see also I’). J) Clones of cells depleted for TAF1B in parallel with panel I, showing reduced clones size and number (green), and competitive cells death at boundaries (magenta, see also J’. Additional data related to this Figure is presented in Figure 2 Supplement 1.

In no case did *Xrp1* mutation eliminate the accumulation of intermediates in *Rp* mutant genotypes (Figure 2B-D; Figure 2 Supplement 1). There were some changes noted in the intermediates that accumulated, however. For example, in *RpS17^+/−^* and *RpS13^+/−^* it seems that more band f accumulates when *Xrp1* is mutated, and less band a. Levels of the pre-rRNA also increase when *Xrp1* is mutated, which can be an indication of elevated rRNA transcription(Sollner-Webb & Tower, 1986), consistent with the faster growth and cell division of *Rp^+/−^ Xrp^+/−^* genotypes than *Rp^+/−^* genotypes(Lee et al., 2018).

In mammalian cells with *Rp* haploinsufficiency, unincorporated 5S RNP, comprising RpL5, RpL11 and the 5S rRNA, activates the transcription factor and tumor suppressor p53 by inhibiting the p53 ubiquitin ligase DM2(Pelava, Schneider, & Watkins, 2016). P53 is responsible for at least some consequences of *Rp* haploinsufficiency in mice, perhaps even including the reduction in translation(Tiu et al., 2020). P53 is also implicated in cell competition in mammals, although not in *Drosophila*, where Xrp1 may acquire some of its functions(Kale, Li, Lee, & Baker, 2015; Baker, Kiparaki, & Khan, 2019). In *Drosophila* it seems that RpS12 is particularly critical for activating Xrp1, through an unknown mechanism(Kale et al., 2018; Boulan, Andersen, Colombani, Boone, & Leopold, 2019; Ji et al., 2019). If a ribosome biogenesis intermediate, for example including RpS12, induced Xrp1 expression, then we predicted that its accumulation and signaling could be prevented by restricting rRNA biogenesis. To test this model, we reduced rRNA synthesis by knockdown of TAF1B, an accessory factor for RNA polymerase I(Knutson & Hahn, 2011). We predicted that Xrp1 expression would be reduced when TAF1B was knocked down in an *Rp*^+/−^ background, and that the knockdown cells would be more competitive than their *Rp*^+/−^ neighbors. Contrary to these predictions, Xrp1 expression was actually higher in *RpS17*^+/−^ dsRNA^TAF1B^ cells than *RpS17*^+/−^ cells(Figure 2E), and *RpS17*^+/−^ dsRNA^TAF1B^ cells underwent cell death at boundaries with *RpS17*^+/−^ territories, suggesting they were less competitive, not more so (Figure 2F). To understand this result, the effect of TAF1B knockdown in otherwise wild type cells was examined, and found to resemble that of *RpS17*^+/−^ dsRNA^TAF1B^ cells. That is, dsRNA^TAF1B^ cells strongly activated Xrp1 expression, and underwent apoptosis at interfaces with wild type cells (Figure 2G,H). This boundary cell death was Xrp1-dependent (Figure 2I,J). Thus, far from rRNA being required for Xrp1 expression and cell competition, as expected if an RNP containing RpS12 activates Xrp1, rRNA depletion appeared to have similar effects to Rp depletion.

It has been suggested that Xrp1 might be released from the nucleolus following nucleolar disruption(Baillon, Germani, Rockel, Hilchenbach, & Basler, 2018). We were unable to detect either resident Xrp1 protein in wild type nucleoli,or altered nucleolar structure in RpS17^+/−^ or RpS18^+/−^ cells mutants by anti-fibrillarin staining (Figure 2-figure supplement 2). It is important to compare Rp^+/−^ and wild type cells at a level where nuclei are present in both, since in mosaic wing discs *Rp^+/−^* nuclei can be displaced basally compared to wild type cells (eg Figure 1-figure supplement 1C,D).

### Reduced protein synthesis is due to PERK-dependent eIF2α phosphorylation in *Rp^+/−^* cells

*Rp* mutations may lead to surplus unused Rp. In yeast, aggregation of unused Rp rapidly affects specific transcription factors, leading to a stress response(Albert et al., 2019; Tye et al., 2019). To explore how Xrp1 reduces translation, if not through reduced ribosome levels, we investigated the phosphorylation of eIF2α, a key mechanism of global regulation of CAP-dependent translation that responds to proteotoxic stress (Hinnebusch & Lorsch, 2012). Strikingly, phosphorylation of eIF2α was increased in a cell-autonomous manner in *Rp^+/−^* cells compared to *Rp^+/+^* cells (RpS3, RpS17, RpS18 and RpL27A were examined) (Figure 3A,B; Figure 3-figure supplement 1A,B). Normal p-eIF2α levels were restored when even one copy of the *Xrp1* gene was mutated, as expected for the Xrp1-dependent process that reduces translation in *Rp^+/−^* cells (Figure 3-figure supplement 1C-E). To verify that eIF2α regulation by Xrp1 was cell-autonomous, we used clonal knockdown with an *Xrp1* dsRNA previously shown to restore normal growth to *Rp^+/−^* cells(Blanco, Cooper, & Baker, 2020). As predicted, *Xrp1* knockdown decreased eIF2α phosphorylation and increased translation rate in a cell-autonomous way (Figure 3E,F), as did knocking-down the gene encoding the Xrp1 heterodimer partner, Irbp18 (Francis et al., 2016; Blanco et al., 2020) (Figure 3-figure supplement 1F, G).

**Figure 3.**
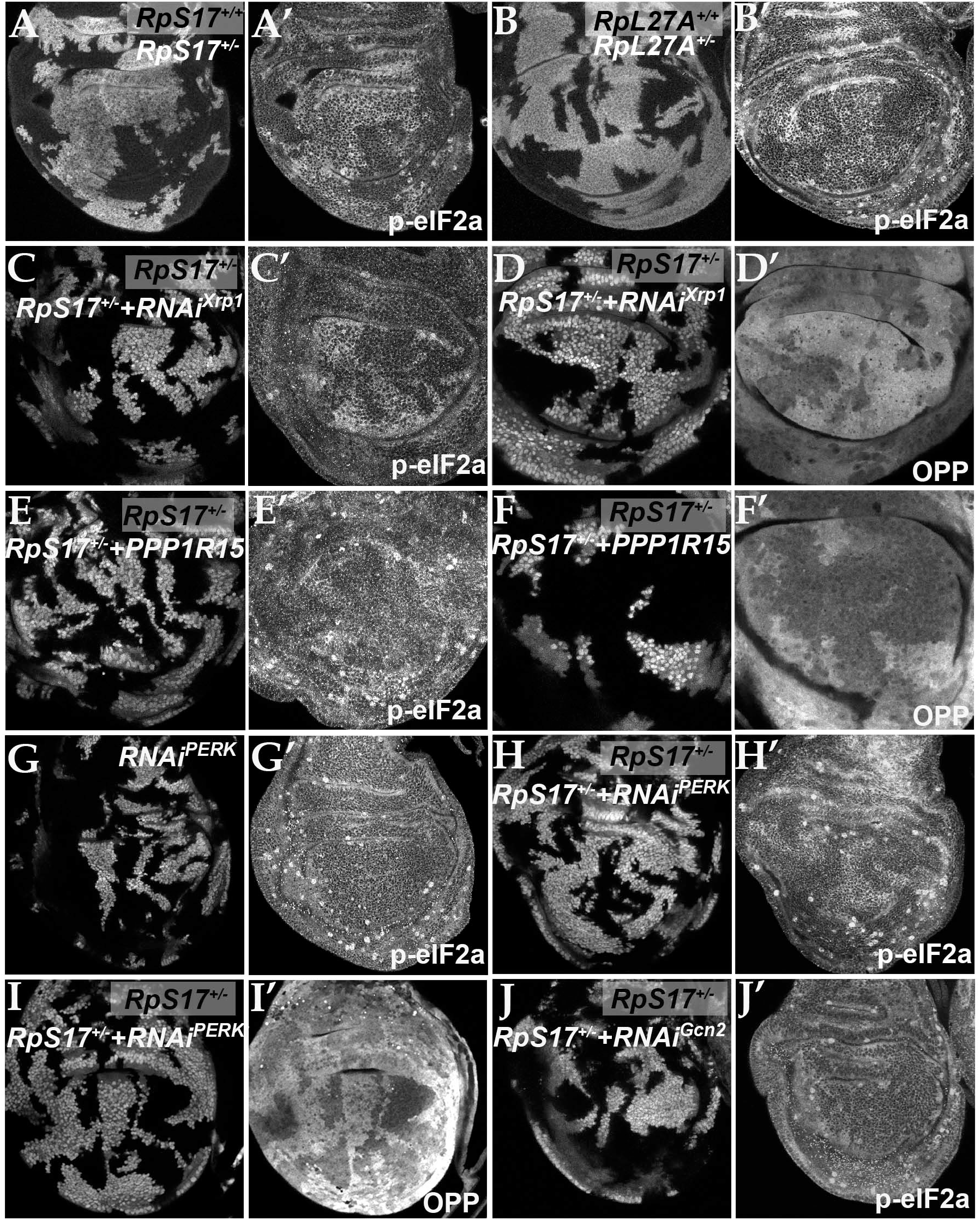
eIF2α is phosphorylated in ribosomal protein mutants via Xrp1 and PERK. Panels A-J show single confocal planes from third instar wing imaginal discs A) Mosaic of *RpS17*^+/−^ and *RpS17*^+/+^ cells. p-eIF2α levels were increased in *RpS17*^+/−^ cells (see A’). B) Mosaic of *RpL27A*^+/−^ and *RpL27A*^+/+^ cells. p-eIF2α levels were increased in *RpL27A*^+/−^ cells (see B’). C) Labelled clones of cells expressing Xrp1-RNAi in a *RpS17*^+/−^ wing disc. p-eIF2α levels were reduced by Xrp1 depletion (see C’). D) Labelled clones of cells expressing Xrp1-RNAi in a *RpS17*^+/−^ wing disc. Translation rate was restored by Xrp1 depletion (see D’). E) Labelled clones of cells over-expressing PPP1R15 in a *RpS17*^+/−^ wing disc,. p-eIF2α levels were reduced by PPP1R15 over-expression (see E’). F) Labelled clones of cells over-expressing PPP1R15 in a *RpS17*^+/−^ wing disc. Translation rate was restored by PPP1R15 over-expression (see F’). G) Labelled clones of cells expressing PERK-RNAi in an otherwise wild type wing disc. p-eIF2α levels were unaffected (see G’). Note that in this and some other panels where mitotic cells are visible near the apical epithelial surface, mitotic figures are labeled by the anti-p-eIF2α antibody, and also lack OPP incorporation. H) Labelled clones of cells expressing PERK-RNAi in a *RpS17*^+/−^ wing disc. p-eIF2α levels were reduced by PERK knockdown (see H’). I) Labelled clones of cells expressing PERK-RNAi in a *RpS17*^+/−^ wing disc. Translation rate was restored by PERK knockdown (see I’). J) Labelled clones of cells expressing Gcn2-RNAi in a *RpS17*^+/−^ wing disc. p-eIF2α levels were not reduced by Gcn2 knockdown (see J’). Further data relevant to this Figure are shown in Figure 3 Supplement 1.

If eIF2α phosphorylation is how Xrp1 reduces translation in *Rp^+/−^* cells, we expected translation to be restored by overexpressed PPP1R15, the *Drosophila* protein homologous to the mammalian p-eIF2α phosphatases, Gadd34 (PPP1R15a) and CReP (PPP1R15b) (Harding et al., 2009; Malzer et al., 2013). Indeed, PPP1R15 cell-autonomously reduced p-eIF2α levels and cell-autonomously restored overall translation levels in multiple *Rp* genotypes, as measured using the click reagent *o*-propargyl puromycin (OPP) (Figures 3E,F; Figure 3-figure supplement 1H,I). These data indicate that it is eIF2α phosphorylation that suppresses translation in *Rp^+/−^* cells.

*Drosophila* contains two potential eIF2α kinases that are thought to respond to particular stresses and not to be activated in unstressed epithelial wing disc cells. When protein kinase R-like endoplasmic reticulum (ER) kinase (PERK), a kinase that phosphorylates eIF2α during the unfolded protein response (Shi et al., 1998; Harding, Zhang, & Ron, 1999; Harding, Zhang, Bertolotti, Zeng, & Ron, 2000; Pakos-Zebrucka et al., 2016), was depleted using RNAi, p-eIF2α levels were unaffected in wild type wing discs (Figure 3G). By contrast, in the *Rp^+/−^* genotypes the levels of p-eIF2α were reduced by PERK depletion (Figure 3H; Figure 3-figure supplement 1J,K). Thus, PERK activity was higher in *Rp^+/−^* cells and responsible for eIF2α phosphorylation there. Consistently, PERK knock-down cell-autonomously restored normal translation levels in multiple *Rp^+/−^* genotypes (Figure 3 I; Figure 3-figure supplement 1L). Depletion of the other eIF2α kinase known in *Drosophila*, Gcn2, did not decrease p-eIF2α levels in *Rp^+/−^* wing disc cells (Figure 3J).

### Xrp1 increases protein aggregation and modified UPR gene expression

Recently, protein aggregates have been detected in the cytoplasm of wing disc cells heterozygous for *RpS3*, *RpS23*, and *RpS26* mutants as foci of ubiquitin or p62 accumulation, reflecting decreased proteasome activity and autophagy(Baumgartner et al., 2021; Recasens-Alvarez et al., 2021). We confirmed the greater accumulation of aggregates in *RpS3^+/−^* and *RpS18^+/−^* cells compared to wild type cells but did not see this for *RpL27A^+/−^* cells(Figure 4A-C). Significantly, another study saw no general increase in autophagy in *RpL14^+/−^* wing discs(Nagata et al., 2019), suggesting this may not occur in mutants affecting the LSU. Importantly, aggregates in *RpS3^+/−^* and *RpS18^+/−^* wing discs were Xrp1-dependent, placing them downstream of Xrp1 activation (Figure 4D-E).

**Figure 4.**
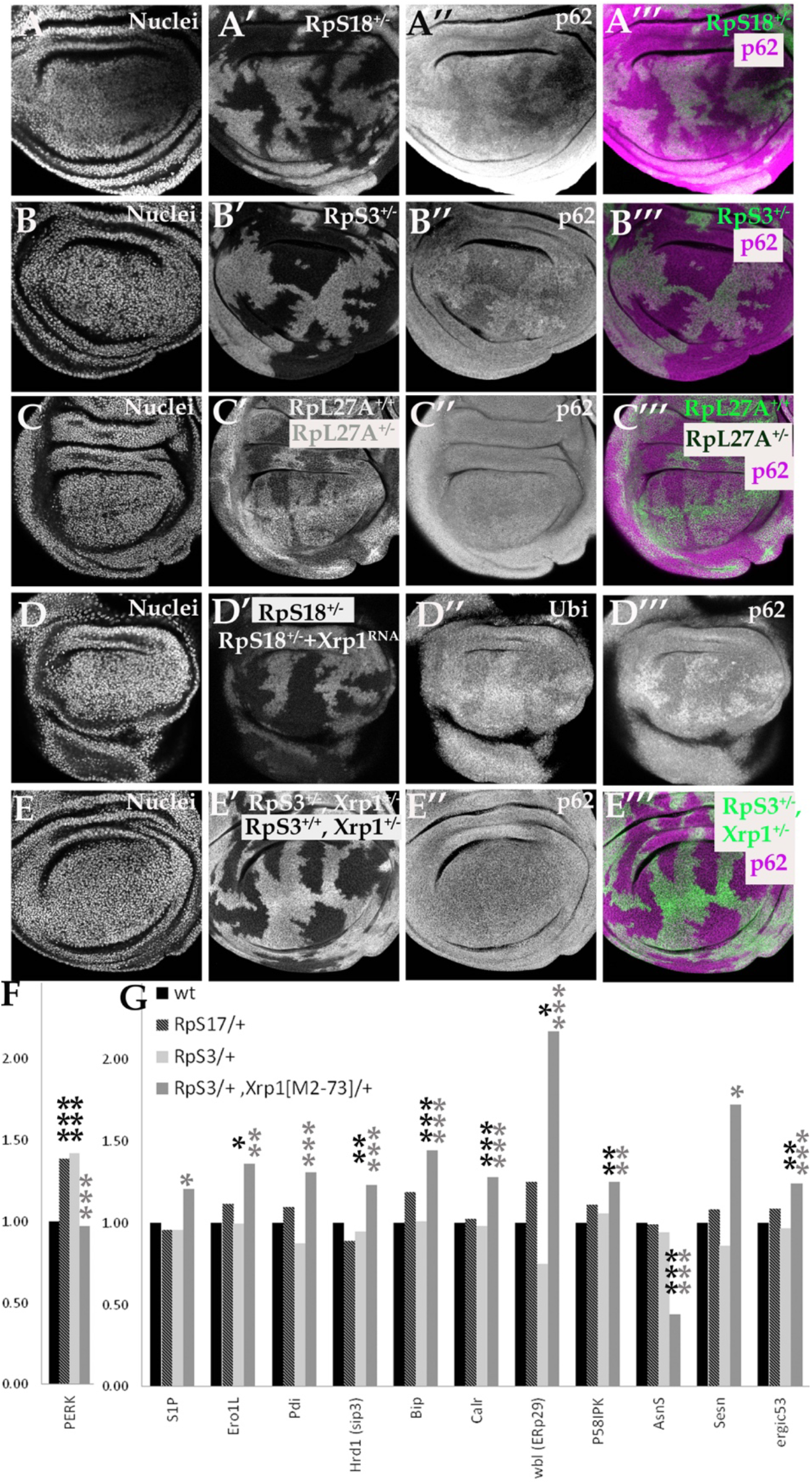
Xrp1-dependent aggregates and gene expression changes in *RpS*^+/−^ cells. Panels A-E show single confocal planes from third instar wing imaginal discs, mosaic for the genotypes indicated. In all cases the plane passes through the central nuclei-containing disk portion for the genotypes shown. A) p62 was higher in *RpS18^+/−^* cells than *RpS18^+/+^* cells. B) p62 was higher in *RpS3^+/−^* cells than *RpS3^+/−^* cells. C) p62 was comparable in *RpL27A^+/−^* cells and *RpL27A^+/+^* cells. D) Labelled clones of cells expressing Xrp1-RNAi in a *RpS18*^+/−^ wing disc. Levels of both p62 and ubiquitinylated proteins were reduced by Xrp1 knock-down. E) Mosaic of *RpS3*^+/−^ and *RpS3*^+/+^ cells in *Xrp1^+/−^* wing disc. No increase in p62 was seen in *RpS3*^+/−^ cells (compare panel B). F) PERK mRNA levels (fold changes in mRNA-seq replicates relative to the wild-type controls according to Deseq2) for the indicated genotypes. PERK mRNA was increased in both *RpS17*^+/−^ and *RpS3*^+/−^ wing discs but not *RpS3*^+/−^, *Xrp1*^M2-73/+^ cells. G) mRNA levels for 11 genes participating in the Unfolded Protein Response. All were significantly affected only in the *RpS3^+/−^ Xrp1^M2-73/+^* genotype. Statistics: Asterisks indicate statistical significance determined by Deseq2 (*: p_adj_<0.05, **: p_adj_<0.005, ***: p_adj_< 0.0005) compared to wild type control (black asterisks) or to *RpS3*^+/−^ genotype (grey asterisks). Comparisons not indicated were not significant ie p_adj_≥0.05 eg *PERK* mRNA in *RpS3^+/−^ Xrp1^M2-73/+^* compared to wild type control. Further data relevant to this Figure are shown in Figure 4 Supplement 1. Data are based on mRNA-sequencing of 3 biological replicates for each genotype.

PERK is a transmembrane protein with a cytoplasmic kinase domain that is a sensor of unfolded proteins within the ER, not within the cytoplasmic or nucleolus(Bertolotti, Zhang, Hendershot, Harding, & Ron, 2000; Harding et al., 2000; Ron & Walter, 2007; Walter & Ron, 2011). Cytoplasmic aggregates can cause unfolded protein accumulation within the ER by competing for proteasomes, however. ER stress also activates Ire-1 and Atf6 in parallel to PERK(Bertolotti et al., 2000; Ron & Walter, 2007; Walter & Ron, 2011; Hetz, 2012; Mitra & Ryoo, 2019). Xbp1-GFP (Sone, Zeng, Larese, & Ryoo, 2013; Mitra & Ryoo, 2019), a reporter for Ire-1 activity, was only inconsistently activated in *Rp^+/−^* wing discs (Figure 4 Figure supplement 1), in agreement with the absence of any transcriptional signature of Atf6 or Xbp1activation in *Rp/+* wing disc mRNA-seq data(Lee et al., 2018). PERK mRNA levels were elevated by 1.4x in both *RpS3^+/−^* and *RpS17^+/−^* wing discs, however (Figure 4F). This increase was statistically very significant, replicated in another group’s data, and entirely dependent on *Xrp1* (Figure 4F)(Kucinski et al., 2017; Lee et al., 2018). *BiP* and 10 other UPR genes were affected differently. Although none were significantly altered in *RpS17^+/−^* or *RpS3^+/−^* discs, all these genes were affected in *RpS3^+/−^ Xrp1^+/−^* wing discs, suggesting that Xrp1 prevents their elevation in *RpS17^+/−^* or *RpS3^+/−^* discs (Figure 4G). Since these genes help restore ER proteostasis (Walter & Ron, 2011), we speculate that Xrp1 might sensitize *Rp^+/−^* cells to PERK activation relative to Atf6 or Xbp1 branches of the UPR(Lin et al., 2007), by elevating the expression of PERK while blunting the usual proteostatic respons. Testing this notion would require manipulating multiple genes in vivo simultaneously.

### eIF2α phosphorylation is sufficient to induce competitive apoptosis, but through Xrp1

We determined whether manipulating p-eIF2α levels alone was sufficient to cause competition of otherwise wild type cells. Consistent with this notion, clones of cells depleted for PPP1R15 were rapidly lost from wing imaginal discs and could rarely be recovered (Figure 5A,B). Under some conditions (longer heat shock) where clones of cells depleted for PPP1R15 survived temporarily, we verified that p-eIF2α was increased and translation reduced compared to nearby wild type cells (Figure 5C,D; Figure 5-figure supplement 1A,B). Such surviving clones were characterized by apoptosis of PPP1R15-depleted cells predominantly at the interface with wild type cells, a sign of cell competition (Figure 5E; Figure 5 figure supplement 1C).

**Figure 5.**
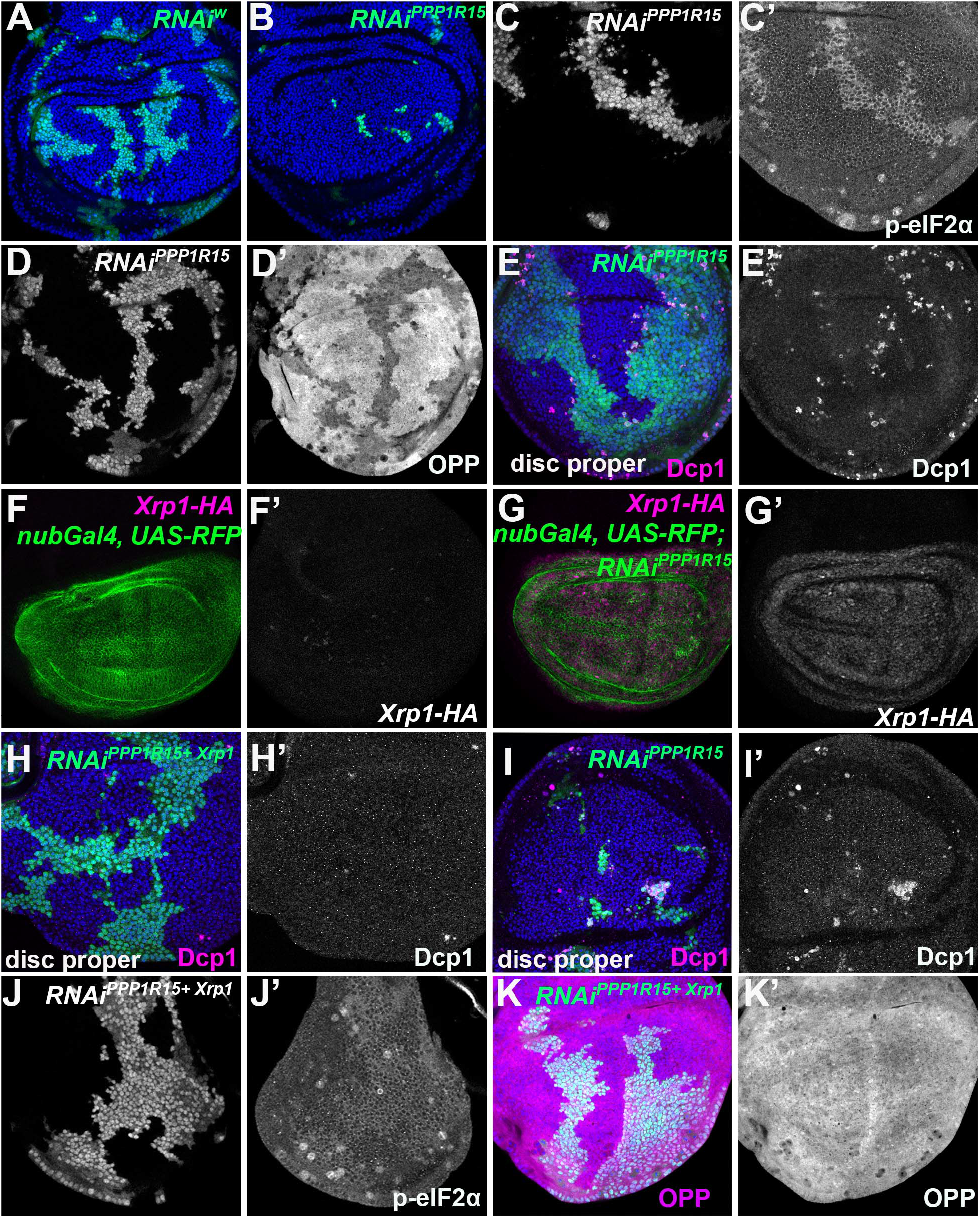
eIF2α phosphorylation induces Xrp1 expression and cell competition. All panels show single confocal planes from third instar wing imaginal discs, mosaic for the genotypes indicated. All the sections pass through the central region of the disc proper containing nuclei in all genotypes, as indicated by the DNA stain in blue in some panels. A) Labelled clones expressing *white* RNAi. Clones induced by 7 min heat shock. B) Labelled clones expressing *PPP1R15* RNAi were fewer and smaller than the control (compare panel A). Clones induced by 7 min heat shock. C) Labelled clones expressing *PPP1R15* RNAi contain phosphorylated eIF2α (see C’). D) Clones induced by 25±525 min heat shock, which results in larger clone areas. Labelled clones expressing *PPP1R15* RNAi reduced translation rate (see D’). E) Labelled clones expressing *PPP1R15* RNAi (green) underwent competitive apoptosis at interfaces with wild type cells (activated caspase Dcp1 in magenta; see also E’). F) *Nub-Gal4* drives gene expression in the wing pouch, shown in green for RFP, with little expression of Xrp1-HA (magenta; see also F’). G) *PPP1R15* RNAi induces Xrp1-HA expression in the wing pouch (magenta; see also G’). H) Labelled clones co-expressing *PPP1R15* RNAi and *Xrp1* RNAi (green) lacked competitive apoptosis (activated caspase Dcp1 in magenta; see also H’). I) Labelled clones expressing *PPP1R15* RNAi (green). Experiment performed in parallel to panel H. Note competitive apoptosis at interfaces with wild type cells (activated caspase Dcp1 in magenta; see also I’), and smaller clone size. Cell death at the basal surface of the same disc shown in Figure 5-Supplement 1F J) Labelled clones co-expressing *PPP1R15* RNAi and *Xrp1* RNAi (green) showed less eIF2α phosphorylation than for *PPP1R15* RNAi alone (compare panel C). Sample prepared in parallel to panel C (in the same tube from fixation to staining). K) *Xrp1* knock-down restored normal translation rate to cell clones expressing *PPP1R15* RNAi (see also K’). Sample prepared in parallel to panel D (in the same tube from fixation to staining). Additional data relevant to this Figure is shown in Figure 5 Supplement 1.

If eIF2α phosphorylation was the downstream effector of Xrp1 that triggers cell competition in *Rp*^+/−^ cells, then PPP1R15 depletion should eliminate cells independently of Xrp1. Like *Rp^+/−^* cells, however, PPP1R15-depleted cells showed strong upregulation of Xrp1 protein (Figure 5F,G; Figure 5 figure supplement 1D). When Xrp1 was knocked-down in PPP1R15-depleted cells, competitive cell death was completely blocked and clone survival improved (Figure H-I; Figure 5-figure supplement 1E-F). Even the p-eIF2α levels in the PPP1R15 depleted clones partially depended on Xrp (compare Figure 5C with Figure 5J), and translation rates were similar to wild type levels in PPP1R15 clones lacking Xrp1 (Figure 5K). Interestingly, PPP1R15 knock-down reduced bristle size, another similarity with *Rp* mutants (Figure 5-figure supplement 1G).

The above data show that eIF2α phosphorylation was sufficient to reduce cell competitiveness in otherwise wild type cells, but only in the presence of Xrp1. It was the mechanism whereby Xrp1 reduced global translation rate in *Rp^+/−^* mutant cells, but was not the downstream effector of Xrp1 for cell competition.

### Interrupting the translation cycle activates Xrp1-dependent cell competition, independently of diminished translation

Phosphorylation of eIF2α inhibits CAP-dependent initiation. To explore further whether reduced translation was sufficient to cause cell competition, we also reduced translation by clonal depletion of translation factors acting(Jackson et al., 2010) at a variety of steps in the translation cycle, not only at initiation but also the 40S-60S subunit joining and elongation steps. Specifically, we depleted eIF4G, eIF5A, eIF6, eEF1α1, and eEF2. eIF4G is part of the eIF4F complex which binds the mRNA 5’cap and recruits SSU to enable translation initiation(Jackson et al., 2010). It is now accepted that eIF5A functions in translation elongation and termination (Saini, Eyler, Green, & Dever, 2009; Schuller, Wu, Dever, Buskirk, & Green, 2017). eEF1α1 delivers aminoacyl-tRNAs to the ribosome and eEF2 also promotes ribosome translocation(Dever & Green, 2012). eIF6 has a role during LSU biogenesis and also in translation initiation(Brina, Miluzio, Ricciardi, & Biffo, 2015).

All these depletions exhibited severe reduction in translation rate in the third instar larvae, as did TAF1B depletion (Figure 6A, E,I,M; Figure 6-figure supplement 1 A,E; the fact that clones of cells expressing these dsRNAs could be recovered with such low translation suggests that translation factor depletion probably exacerbates over time, initially being insufficient to prevent translation and growth, but eventually becoming severe). Importantly, all of these translation factor depletions resulted in dramatic induction of apoptosis in depleted cells that were close to wild type cells, suggesting that differences in translation rate might be sufficient to initiate cell competition (Figure 6B,F,J; Figure 6-figure supplement 1 B,F). Interestingly, in all these cases translation increased in wild type cells near to the affected clones, something that was rare adjacent to *Rp*^+/−^ cells and not seen adjacent to cells depleted for PPP1R15, although it was observed near to TAF1B depleted cells (Figure 6A,E,I,M; Figure 6-figure supplement 1 A,E). Phosphorylated RpS6 accumulated in wild type cells adjacent to TAF1B depleted cells, suggesting that a non-autonomous activation of Tor accounts for the increased translation in cells nearby those with translation deficits (Figure 6N)(Laplante & Sabatini, 2012; Romero-Pozuelo, Demetriades, Schroeder, & Teleman, 2017).

**Figure 6.**
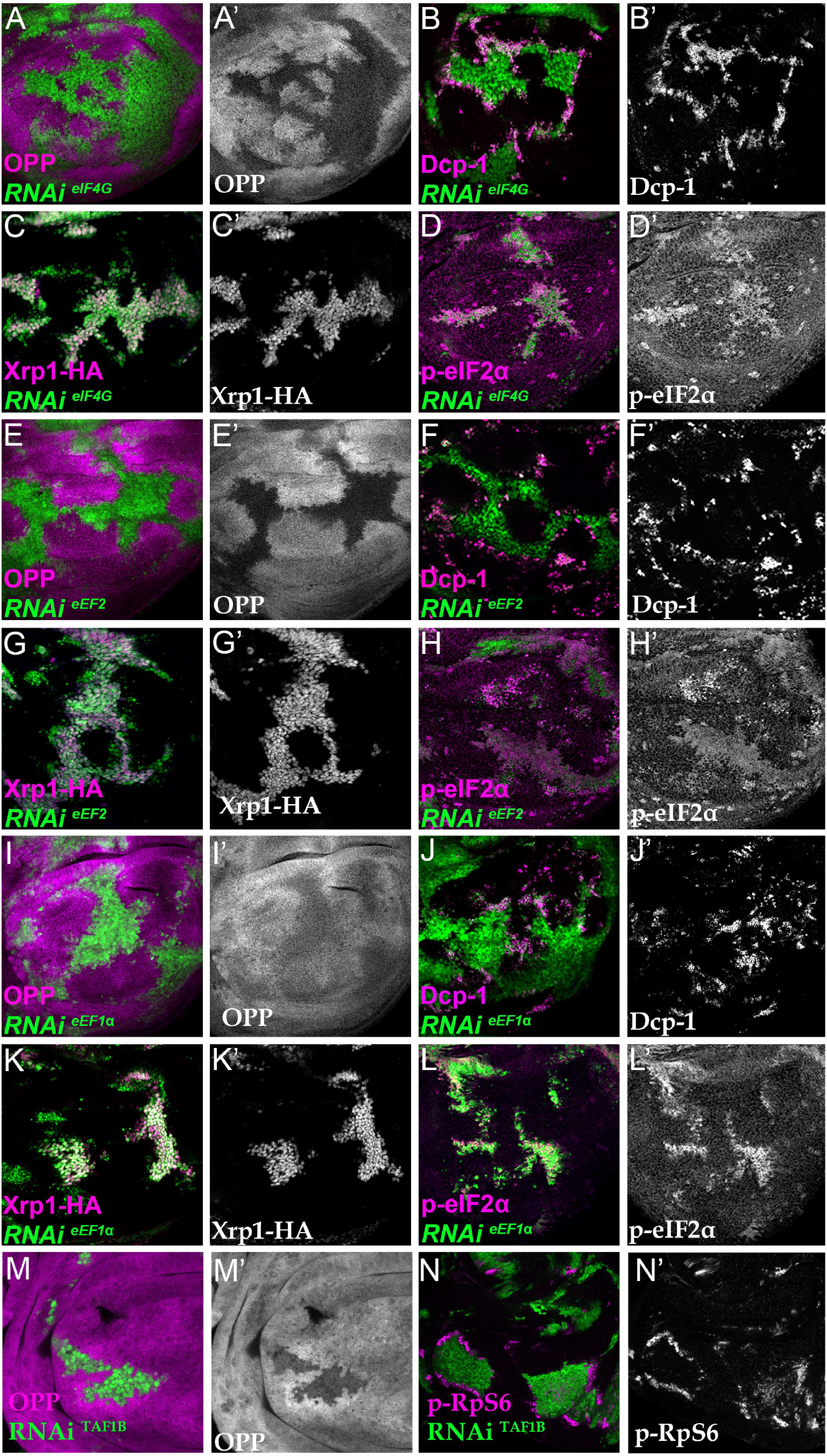
Depletion of translation factors induces Xrp1 expression, eIF2α phosphorylation, reduced translation, and cell competition. Clones of cells depleted for translation factors are labelled in green. In each case, translation factor depletion reduced translation rate, resulted in competitive cell death at interfaces with wild type cells, induced Xrp1-HA expression, and led to eIF2α phosphorylation. Translation rate, dying cells (activated caspase Dcp1), Xrp1-HA and p-eIF2α are indicated in magenta and in separate channels as labelled. A-D) Clones expressing RNAi for eIF4G. E-H) Clones expressing RNAi for eEF2. I-L) Clones expressing RNAi for eEF1α. In all cases (panels A,E,I), wild type cells near to cells depleted for translation factors show higher translation rate than other wild type cells. M) Clones of cells depleted for TAF1B (green) also showed a cell-autonomous reduction in translation rate and non-autonomous increase in nearby wild type cells (translation rate in magenta, see also M’). N) Clones of cells depleted for TAF1B (green) showed a non-autonomous increase in RpS6 phosphorylation in nearby cells (magenta, see also N’). Additional data relevant to this Figure is shown in Figure 6 Supplement 1 and Figure 6 Supplement 2.

To confirm that translation factor depletion affected translation directly, and downstream of Xrp1 and PERK, Xrp1 expression and eIF2α phosphorylation were examined. Unexpectedly, depletion for translation factors was associated with both cell-autonomous induction of Xrp1 expression and eIF2α phosphorylation (Figure 6C,D,G,H,K,L; Figure 6 Figure Supplement 1C,D,G,H; Figure 6-figure supplement 2). The levels were at least comparable to those of TAF1B-depleted cells (Figure 6I,J). When Xrp1 was knocked-down, PPP1R15 overexpressed, or PERK depleted simultaneously with translation factor depletion, the translation factor depletions behaved similarly to one another. PPP1R15 overexpression was sufficient to reduce eIF2α phosphorylation to near or even below control levels (Figure 7A,D,G), but this did not restore normal translation rates (Figure 7B,E,H). There was no rescue of competitive cell death (Figure 7C,F,I; Figure 7-figure supplement 1A,C) or Xrp1 expression (Figure 7J-L; Figure 7-figure supplement 1 B,D). PERK knock-down similarly did not affect Xrp1 expression or rescue competitive cell death in translation-factor knock-downs (Figure 7 – figure supplement 2). Knockdown of Xrp1 reduced levels of eIF2α phosphorylation in some cases (Figure 7M,P Figure 7-figure supplement 1E), although for eIF5A and eEF1α1 the reduction was only partial so that both the eIF5A Xrp1 depleted and eEF1α1 Xrp1 depleted cells retained more eIF2α phosphorylation than wild type cells(Figure 7S; Figure 7-figure supplement 1E). For all the translation factors, however, Xrp1 depletion eliminated cell death at the competing cell boundaries, irrespective of whether eIF2α phosphorylation remained (Figure 7O,R,U; Figure 7-figure supplement 1G,J). We also found that overall translation rate, as estimated by OPP incorporation, was only partially restored by simultaneous Xrp1 depletion from most translation factor knock-down cells, and remained lower than wild type cells (Figure 7N,Q; Figure 7-figure supplement 1C 7B,E, Figure 7-figure supplement 1F). Two exceptions were eIF6 and eEF1α1. Remarkably, simultaneous knock-down of Xrp1 along with either of these genes resulted in translation rates similar to or higher than in wild type cells (Figure 7T; Figure 7-figure supplement E).

**Figure 7.**
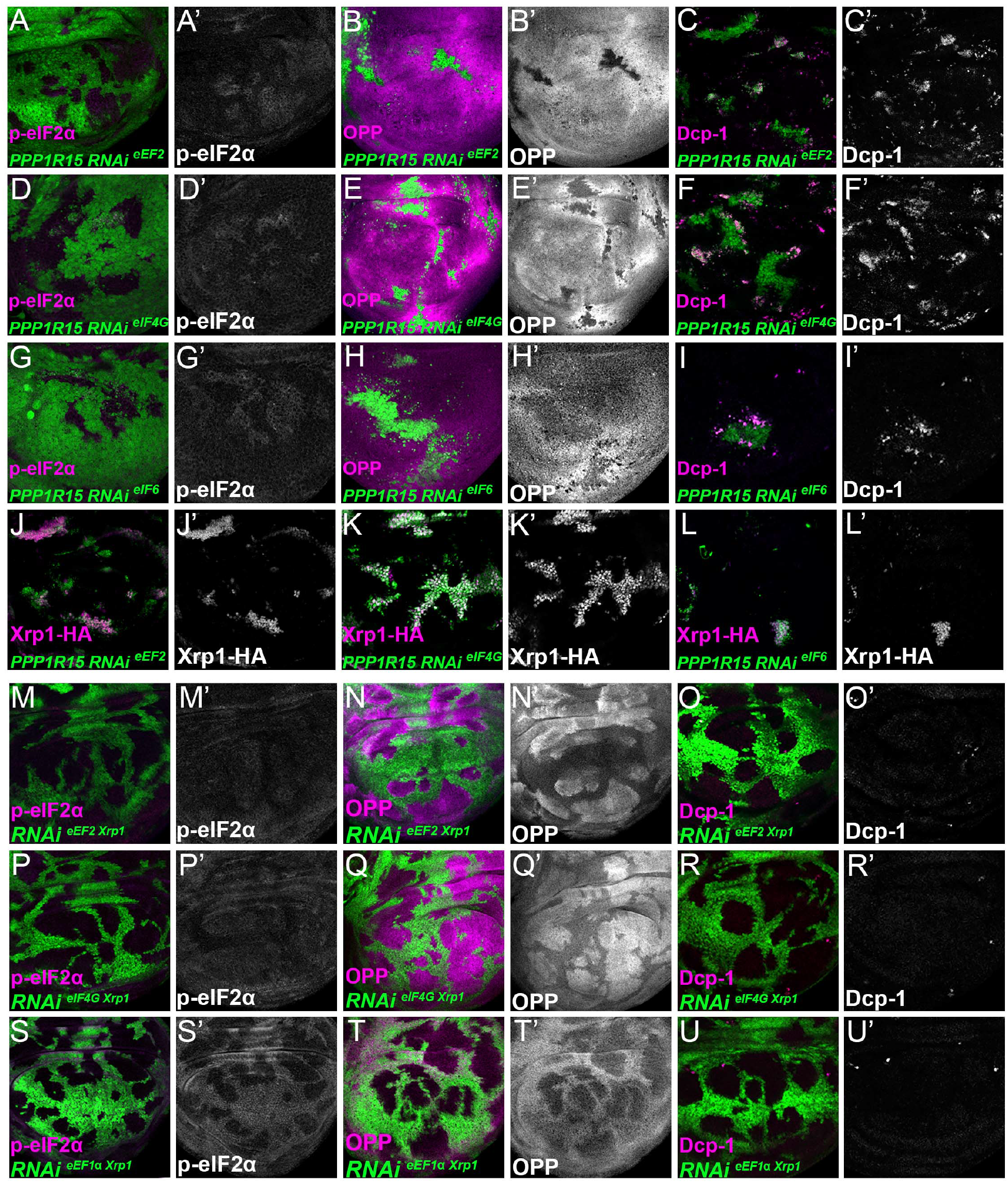
Interrupting the translation cycle activates Xrp1-dependent cell competition, independently of diminished translation. Single confocal planes from third instar wing imaginal discs. p-eIF2α levels, translation rate (ortho-propargyl puromycin), dying cells (activated caspase Dcp1) and Xrp1-HA are indicated in magenta and in separate channels as labelled. (A-L) Clones of cells depleted for translation factors which also overexpress PPP1R15 are shown in green. In each case, PPP1R15 overexpression was sufficient to reduce eIF2α phosphorylation to near control levels (or even lower), but it did not restore normal translation rates, did not affect Xrp1-HA levels and did not reduce competitive cell death. A-C) Clones co-expressing PPP1R15 and RNAi for eEF2. D-F) Clones co-expressing PPP1R15 and RNAi eIF4G. G-I) Clones co-expressing PPP1R15 and RNAi for eIF6. J-K) Xrp1-HA expression (magenta) in clones co-expressing PPP1R15 and RNAi for eEF2 (J), eIF4G (K), or eIF6 (L). (M-U) Clones of cells depleted for translation factors which also express Xrp1-RNAi are shown in green. (M-O) Clones depleted for Xrp1 as well as eEF2 expressed phospho-eIF2α at near to control levels, only partially restored overall translation rate, but lacked competitive cell death. (P-R) Clones depleted for Xrp1 as well as eIF4G expressed phospho-eIF2α at near to control levels, only partially restored overall translation rate, but lacked competitive cell death. (S-U) Clones depleted for Xrp1 as well as eEF1α1 retained high levels of eIF2α phosphorylation but actually showed a global translation rate higher than wild type cells. They lacked competitive cell death.

These results unexpectedly show that translation factor depletion triggers similar effects to depletion of ribosome components, in which Xrp1 expression leads to eIF2α phosphorylation and to cell competition. The results separate eIF2α phosphorylation from cell competition, however, because Xrp1-dependent cell competition continued even when eIF2α phosphorylation levels was restored to normal by PPP1R15 overexpression, and because remaining eIF2α phosphorylation in eIF5A Xrp1-depleted and eEF1α1 Xrp1-depleted cells did not lead to cell competition. The results also separate cell competition from differences in translation levels, because no competitive cell death was observed in eIF4G Xrp1-depleted, eIF5A Xrp1-depleted, and eEF2 Xrp1-depleted cells, even though their translation was lower than the nearby wild type cells. Indeed, depletion for eIF6 or eEF1α1 induced Xrp1 and cell competition, even though without Xrp1 these cells seemed to translate at similar or higher rates to their neighbors. These results focus attention on Xrp1 as the key effector of cell competition, irrespective of eIF2α phosphorylation and overall translation rate.

These results also raise the question of whether *Rp* haploinsufficiency, rRNA depletion, eIF2α phosphorylation, and translation factor depletion all activate Xrp1 through a common pathway. In *Rp*^+/−^, Xrp1 expression genotypes depends on a specific ribosomal protein, RpS12, and is almost completely prevented by *rpS12^G97D^*, a missense allele that specifically affects this aspect of RpS12 function(Lee et al., 2018; Ji et al., 2019). We found that *rpS12^G97D^* homozygosity also reduced Xrp1 induction when TAF1B was depleted (Figure 8A-C), but had much less effect when eEF2 was depleted (Figure 8D-E). Thus, the mechanism of Xrp1 activation may resemble that in *Rp*^+/−^ cells when rRNA synthesis is affected, but appears distinct when translation factors are inhibited.

**Figure 8.**
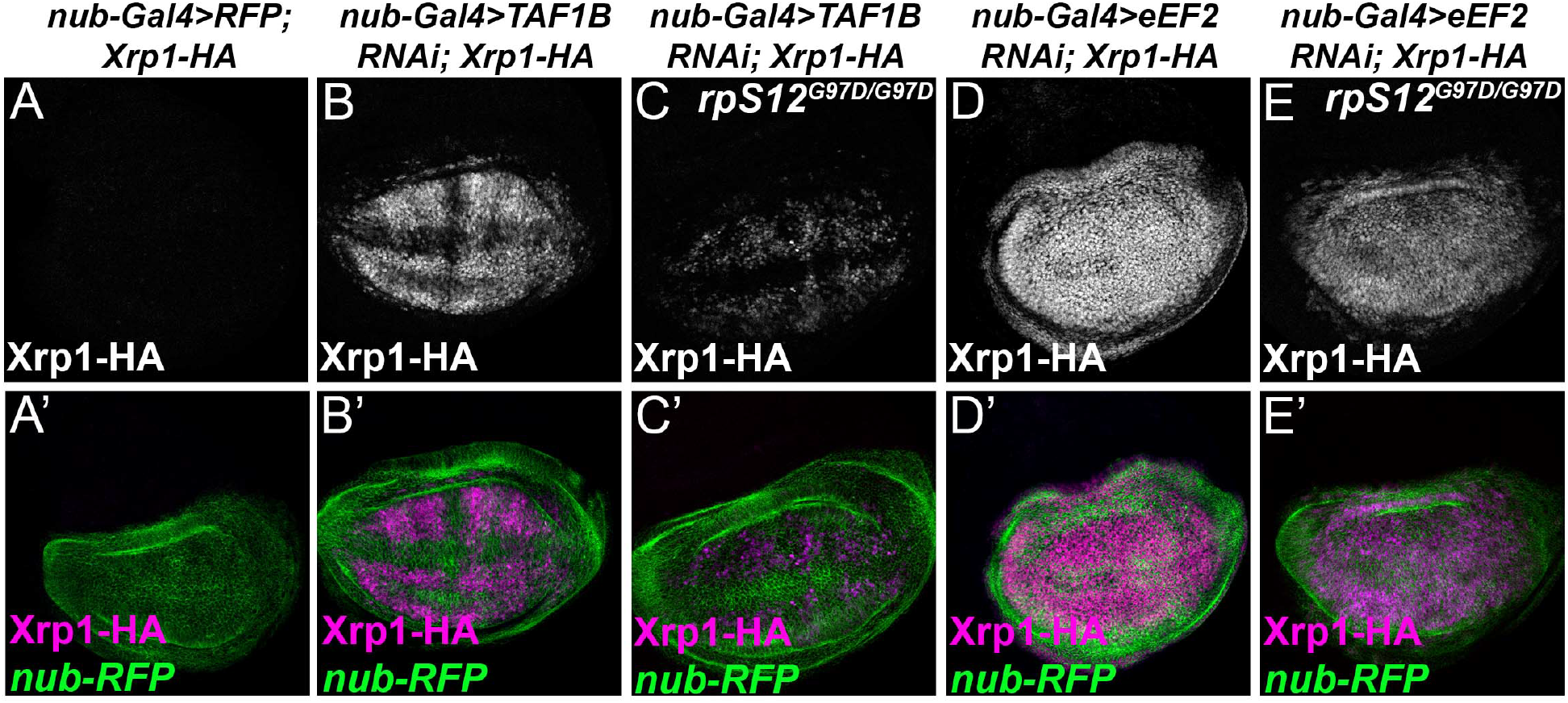
RpS12-dependence of Xrp1 expression. Figures show projections of Xrp1-HA expression from the wing discs of indicated genotypes. A) Neglegible Xrp1-HA (magenta in A’) was expressed in control discs where *nub-Gal4* drove only reporter RFP expression in the wing pouch (green in A’-E’). TAF1B knockdown resulted in Xrp1-HA expression (magenta in B’). C) Xrp1-HA expression was greatly reduced when TAF-1B was knocked-down in the *rpS12^G97D^* background (see also magenta in C’). D) eEF2 knockdown resulted in strong Xrp1-HA expression (magenta in D’). E) Xrp1-HA expression was only moderately reduced when eEF2 was knocked-down in the *rpS12^G97D^* background (see also magenta in E’).

### Xrp1 is a transcription factor that regulates cell competition

Xrp1 as a key mediator of multiple defects in ribosome biogenesis or function. Xrp1 is a sequence-specific DNA-binding protein implicated in genome maintenance, and binds directly to sequences of the P element whose transposition it promotes(Akdemir, Christich, Sogame, Chapo, & Abrams, 2007; Francis et al., 2016). Xrp1 also controls expression of many genes at the mRNA level(Lee et al., 2018), and other similar bZip proteins are transcription factors(Tsukada, Yoshida, Kominato, & Auron, 2011).

To test whether Xrp1 is a transcription factor, we used a dual-luciferase reporter system in transfected S2 cells (Figure 9A-D; Figure 9 Supplement 1). Luciferase reporter plasmids were either based on the widely-used core promoter of the Drosophila hsp70 gene, or on a 400bp genomic sequence spanning the transcription start site of the *Xrp1* gene itself (Figure 9-figure supplement 2). We cloned 8x repeats of either of two different matches to the 10bp Xrp1/Irbp18 consensus binding site in vitro(Zhu et al., 2011), which is similar to that recently deduced from ChIP-Seq following Xrp1 overexpression in vivo(Baillon et al., 2018) (Target 1 and Target 3) or of the sequence footprinted by Xrp1/Irbp18 on the P element terminal repeat (Francis et al., 2016)(target 2), which also contains a consensus match (Figure 9A,B). When Xrp1 expression was induced in transfected S2 cells, each of these Xrp1-binding sequences conferred 3x-8x activation of luciferase expression, whereas scrambled sequences were inactive (Figure 9C,D, Figure 9-figure supplement 1A,B). In the case of target 1, several-fold further induction was achieved by co-transfection and induction of Irbp18 expression, culminating in 23x stimulation of luciferase expression by repeats of the Target 1 sequence in conjunction with the hsp70 basal promoter (Figure 9-figure supplement 1A). Irbp18 alone was inactive in the absence of transfected Xrp1(Figure 9C,D; Figure 9-figure supplement 1A,B). Thus, the Xrp1/Irbp18 heterodimer stimulated transcription through its cognate binding sequences.

**Figure 9.**
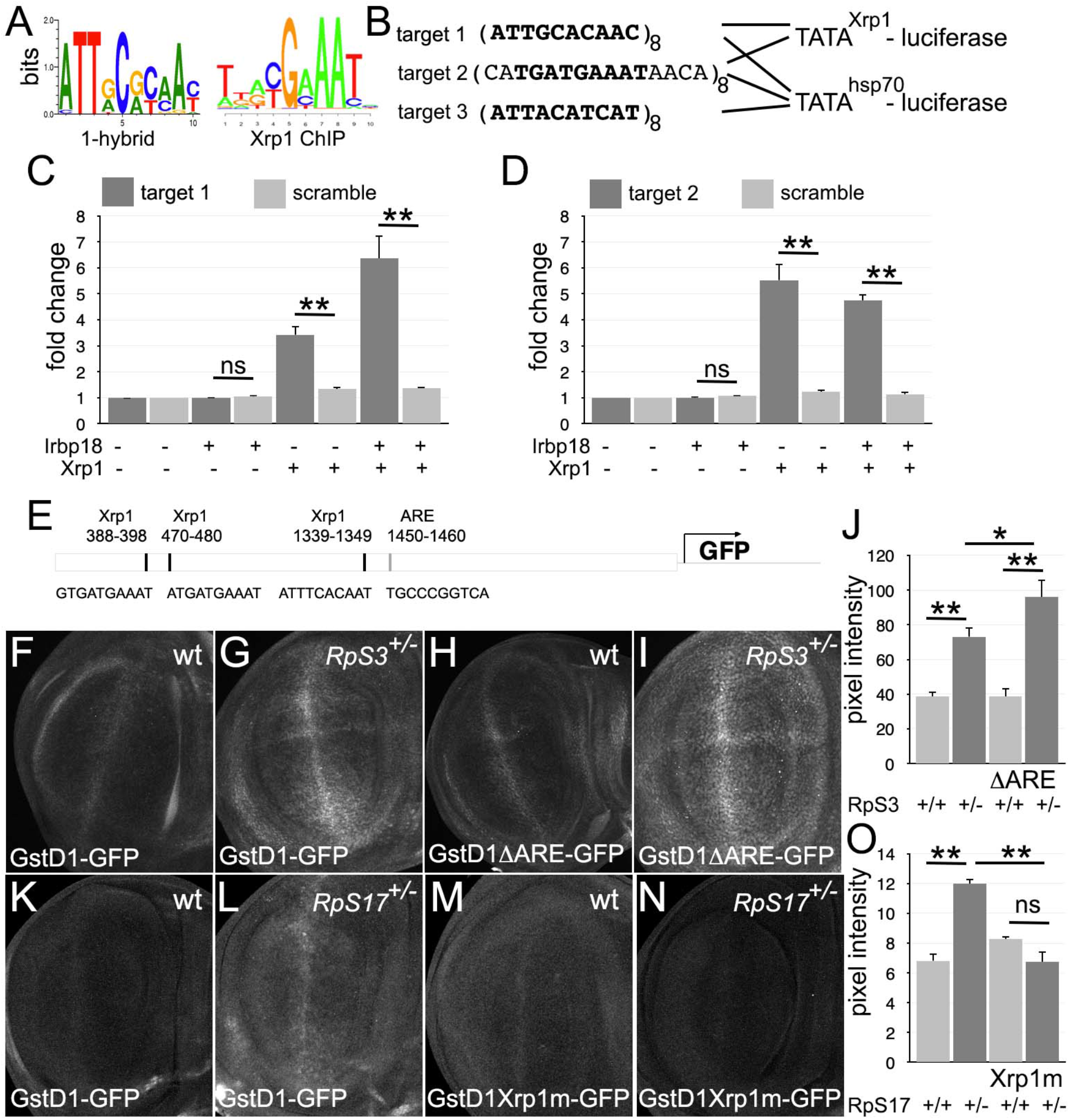
Transcriptional regulation by Xrp1. A) Similar consensus binding site of Xrp2/Irbp18 defined by bacterial 1-hybrid studies(Zhu et al., 2011) and by Xrp1 ChIP from *Drosophila* eye imaginal discs overexpressing an Xrp1-HA protein(Baillon et al., 2018). B) Xrp1 binding motif sequences multimerized in luciferase reporter plasmids upstream of transcription start sites from the *Xrp1* gene or from the *hsp70* gene. Targets 1 and 3 were based on the 1-hybrid consensus, target 2 is the P element sequence footprinted by Xrp1-Irbp18(Francis et al., 2016). The match to the consensus sites is shown in bold type. C) Luciferase assays following transfection of reporters and protein expression plasmids into S2 cells. The target 1-TATA^Xrp1^ reporter showed sequence-specific activation by transfected Xrp1. Transfected Irb18 alone had no effect, but synergized with Xrp1. Exact p-values for comparisons between target 1 reporters and scrambled reporters were: Padj=6.16, Padj=0.00827, Padj=3.47×10^−7^ respectively. D) Luciferase assays following transfection of reporters and protein expression plasmids into S2 cells. The target 2-TATA^Xrp1^ reporter showed sequence-specific activation by transfected Xrp1. Transfected Irbp18 alone had no effect. Exact p-values for comparisons between target 2 reporters and scrambled reporters were: Padj=4.21, Padj=2.00×10^−8^, Padj=1.96×10^−7^ respectively. E) Potential regulatory sequences in the 2.7kb upstream intergenic fragment used in the GstD1-GFP reporter(Brown et al.; Sykiotis & Bohmann, 2008). 3 Xrp1-binding motifs and the antioxidant response element (ARE) are indicated. F-I) and K-N) show projections from the central disc-proper regions of wing discs expressing reporter transgenes in the indicated genetic backgrounds. F) baseline GstD1-GFP expression in the wild type wing disc. G) Elevated GstD1-GFP expression in the *RpS3^+/−^* wing disc. H) baseline GstD1ΔARE-GFP expression in the wild type wing disc. I) Elevated GstD1ΔARE-GFP expression in the *RpS3^+/−^* wing disc. J) Quantification of these results. Average pixel intensity from wing pouch regions was measured. Mean ± SEM from multiple samples is shown. N=5 for each genotype. Exact P values were: for GstD1-GFP in *RpS3^+/−^* compared to *RpS3^+/+^*, Padj=0.00257; for GstD1ΔARE-GFP in *RpS3^+/−^* compared to *RpS3^+/+^*, Padj=2.55×10^−5^; for GstD1-GFP in *RpS3^+/+^* compared to GstD1ΔARE-GFP in *RpS3^+/+^*, Padj=0.993; for GstD1-GFP in *RpS3^+/−^* compared to GstD1ΔARE-GFP in *RpS3^+/−^*, Padj=0.0313. K) baseline GstD1-GFP expression in the wild type wing disc. L) Elevated GstD1-GFP expression in the *RpS17^+/−^* wing disc. M) baseline expression of GstD1-GFP with all 3 Xrp1-binding motifs mutated in the wild type wing disc. N) Expression of GstD1-GFP with all 3 Xrp1-binding motifs mutated was similar in the *RpS17^+/−^* wing disc to the wild type control. O) Quantification of these results. Average pixel intensity from wing pouch regions was measured. Mean ± SEM from multiple samples is shown. N=5,6,5,6 for respective samples. Exact P values were: for GstD1-GFP in *RpS3^+/−^* compared to *RpS3^+/+^*, Padj=2.34×10^−6^; for GstD1mXrp1-GFP in *RpS3^+/−^* compared to *RpS3^+/+^*, Padj=0.116; for GstD1-GFP in *RpS3^+/+^* compared to GstD1mXrp1-GFP in *RpS3^+/+^*, Padj=0.112; for GstD1-GFP in *RpS3^+/−^* compared to GstD1mXrp1-GFP in *RpS3^+/−^*, Padj=1.19×10^−6^. K) baseline GstD1-GFP expression in the wild type wing disc. Statistics: 1-way ANOVA with Bonferroni-Holm correction for multiple testing was performed for the data shown in panels C,D,J,O. Data in panel C,D were based on triplicate measurements from each of 3 biological replicates for each transfection.

It has been suggested that an oxidative stress response in *Rp*^+/−^ cells leads to competition with wild type cells(Kucinski et al., 2017; Baumgartner et al., 2021). *Rp*^+/−^ cells express GstD1 reporters, whose transcription is activated by Nrf2, the master regulator of oxidative stress responses(Kucinski et al., 2017). Because the genes expressed in *Rp*^+/−^ cells are also enriched for Xrp1 binding motifs, some of these genes might be activated directly by Xrp1, including GstD1(Ji et al., 2019). The GstD1-GFP reporter used to report oxidative stress in *Rp*^+/−^ cells contains a 2.7 kb genomic fragment that contains an Antioxidant Response Element (ARE) bound by the Nrf2/Keap1 dimer at position 1450-1460(Figure 9E). Deletion of this motif abolishes GstD1-GFP induction in response to oxidative stress(Sykiotis & Bohmann, 2008). Recently, Brown et al identified Xrp1 binding motifs within the same GstD1-GFP reporter, and showed that these sequences are required for Xrp1-dependent induction in response to ER stress(Brown, Mitra, Roach, Vasudevan, & Ryoo). We therefore compared induction of GstD1-GFP reporters in *Rp*^+/−^ wing discs where the reporter sequences were either wild type, deleted for the Nrf2 binding motif, or mutated at the Xrp1-binding motifs (Figure 9E). We found that the Nrf2 binding motif was dispensable for GstD1-GFP induction in *Rp*^+/−^ wing discs, whereas the Xrp1 sites were required, consistent with induction of GstD1-GFP and perhaps other genes as direct transcriptional targets of Xrp1, not Nrf2(Figure 9F-O).

## DISCUSSION

We explored the mechanisms by which *Rp* mutations affect *Drosophila* imaginal disc cells, causing reduced translation and elimination by competition with wild type cells in mosaics. Our findings reinforced the key role played by the AT-hook bZip protein Xrp1, which we showed is a sequence-specific transcription factor responsible for multiple aspects of not only the *Rp* phenotype, but also other ribosomal stresses (Figure 10). It was Xrp1, rather than the reduced levels of ribosomal subunits, that affected overall translation rate, primarily through PERK-dependent phosphorylation of eIF2α. Phosphorylation of eIF2α, as well as other disruptions to ribosome biogenesis and function such as reduction in rRNA synthesis or depletion of translation factors, were all sufficient to cause cell competition with nearby wild type cells, but this occurred because all these perturbations activated Xrp1, not because differences in translation levels between cells cause cell competition directly (Figure 10). Other features of *Rp^+/−^* cells, including protein aggregation and activation of ‘oxidative stress’ genes, were also coordinated by Xrp1, contrary to the notion that proteotoxic stress directly triggers oxidative damage in *Rp^+/−^* cells. These findings confirm the central importance of the transcriptional response to *Rp* mutations, and to other disruptions of ribosome biogenesis and function. They suggest therapeutic approaches to ribosomopathies, and have implications for the surveillance of aneuploid cells.

**Figure 10.**
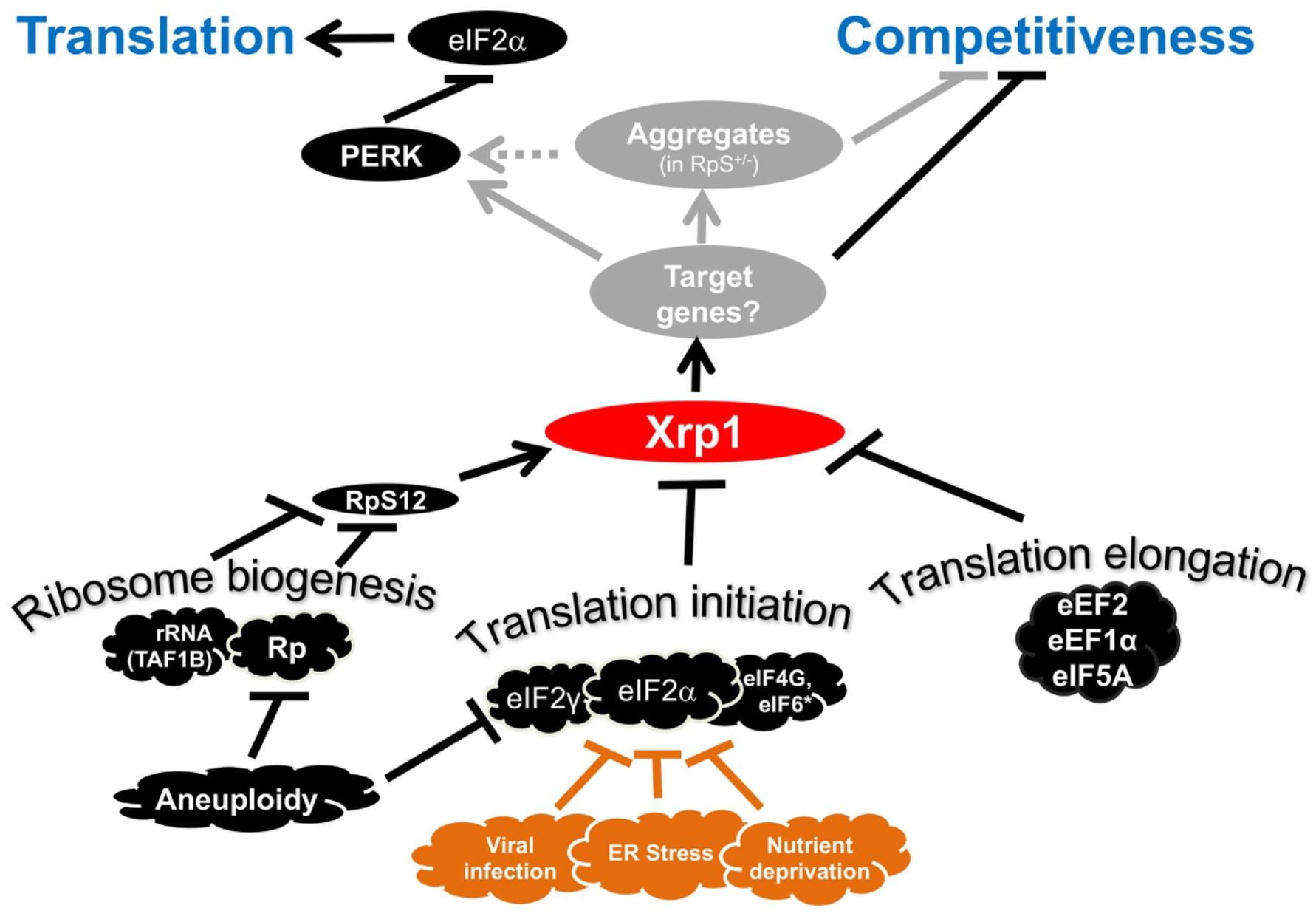
Transcriptional responses to Ribosome defects. Multiple consequences of defects in ribosome biogenesis, translation initiation, and translation elongation, depend on the transcription factor Xrp1 in imaginal disc cells. Xrp1 is responsible for, or contributes to, reduced translation in response to these defects, through the PERK-dependent phosphorylation of eIF2α, a global regulator of CAP-dependent translation initiation. Xrp1 protein expression also marks imaginal disc cells for elimination in competition with wild type cells. Cell competition often correlates with differences in translation rate because so many ribosome stresses activate Xrp1. This includes reduced eIF2 activity, as caused by eIF2α phosphorylation, or eIF2γ haploinsufficiency, but these are not sufficient to trigger cell competition without Xrp1. We speculate that other cellular stresses that phosphorylate eIF2α, including ER stress, nutrient deprivation, or (in mammals) infection with certain viruses might mark cells for competition, or interfere with cell competition that recognizes aneuploid cells on the basis of *Rp* or eIF2γ gene haploinsufficiency. It is notable that defective Tor signaling, which also reduces global translation rate, does not cause cell competition, (Baumgartner et al., 2021), making the molecular mechanism of Xrp1 induction uncertain, although several genetic pathways have been shown to induce Xrp1, including dependence on RpS12 in *Rp^+/−^* cells and TAF1B-depleted cells(Akdemir et al., 2007; Chapin et al., 2014; Lee et al., 2018; Ji et al., 2019).

### Ribosome levels are modestly affected by *Rp* gene haploinsufficiency

Multiple assays show that ribosome subunit concentration is only moderately affected by *Rp* haploinsufficiency. In *RpL* mutants, we have seen 15-20% reduction in LSU concentrations, and 0-25% reduction in SSU concentrations, whereas in *RpS* mutants we have seen 20-25% reduction in SSU concentrations and 0-10% increase in LSU concentrations. Consistent with this, mass spec measurements of *RpS3*^+/−^ and *RpS23*^+/−^ *Drosophila* wing discs found that other RpS proteins were typically under-represented but RpL subunits over-represented in these genotypes(Baumgartner et al., 2021; Recasens-Alvarez et al., 2021). Broadly similar results have been reported in yeast, and the major changes in yeast gene expression following reduced Rp expression also seem to have a transcriptional basis, not translational(Cheng et al., 2019). We found that ribosomal subunit levels were unaffected by *Xrp1*, suggesting that changes to their levels are likely more direct consequences of the *Rp* mutations.

The fact that ribosome subunit concentrations change modestly, and differently between mutations affecting LSU and SSU proteins, does not rule out functional consequences of these changes, which could depend more on the concentrations of free SSU and LSU than on their total concentrations. It suggests, however, that cellular and animal models of DBA that have generally sought to achieve a 50% reduction in Rp protein expression(Heijnen et al., 2014; Khajuria et al., 2018) could be significantly more severe than occurs in DBA patients, and that actual ribosome subunit concentrations should be measured in DBA patient cells to guide future models.

Multiple explanations for the modest effects of *Rp* haploinsufficiency on ribosome subunit number are possible. We particularly point out that, even if expression of a particular Rp is reduced in proportion to a 50% reduction in mRNA level, the respective protein <u>concentration</u> (ie number of molecules/cell volume) is unlikely to fall to 50%, because ribosomes are required for cellular growth, so that an *Rp* mutation affects the denominator in the concentration equation, as well as the numerator. It is even possible that a 50% reduction in its rate of Rp synthesis could leave steady state ribosome subunit concentration unaffected, if cellular growth rate was slowed by the same amount.

### *Rp* mutant cells accumulate ribosome biogenesis intermediates but protein aggregates requires Xrp1

The other proximate effect of *Rp* mutations is the accumulation and disposal of ribosome components that are left unused. We confirmed that ribosome assembly intermediates do indeed accumulate in *Drosophila* wing discs following *Rp* haploinsufficiency. Reduction in polI activity parallel with *Rp* haploinsufficiency did not suppress the *Rp* phenotype, however, providing no support for a signaling species containing both RNA and unused Rp. In yeast, aggregates of unused Rp rapidly trigger specific transcriptional responses(Albert et al., 2019; Tye et al., 2019). In *Drosophila Rp*^+/−^ cells, Xrp1 expression depends particularly on RpS12, rather than on all unused Rp(Lee et al., 2018; Boulan et al., 2019; Ji et al., 2019). Significantly, the protein aggregates that had been detected in *Rp* mutant *Drosophila* wing discs (Baumgartner et al., 2021; Recasens-Alvarez et al., 2021) appeared specific for mutations in SSU proteins, and were a downstream consequence of Xrp1 activity rather than direct consequence of *Rp* mutations (Figure 10). Reduced translation and cell competition are features of both RpS and RpL mutants(Lee et al., 2018). On the other hand, differences between effects of RpS and RpL mutations, which are also seen in yeast (Cheng et al., 2019), could account for the contradictory findings that autophagy seems protective for cells mutated for RpS genes(Baumgartner et al., 2021; Recasens-Alvarez et al., 2021), but promotes competitive apoptosis in cells mutated for an LSU gene(Nagata et al., 2019). Although it seems clear that unused Rp aggregate in yeast, whether the particular protein aggregates visible in *Drosophila* cells contain Rp or represent the primary feature of ‘loser status’ in cell competition requires more investigation. Alternatively, *Rp* mutations may cause rapid transcriptional reprogramming in *Drosophila* cells(Lee et al., 2018), as also occurs in yeast(Albert et al., 2019; Cheng et al., 2019; Tye et al., 2019).

### *Rp* mutants affect global translation rate through eIF2α

The main mechanism by which Xrp1 suppresses global translation in *Rp^+/−^* mutants was shown to be PERK-dependent phosphorylation of eIF2α. PERK is activated by ER stress, although the IRE/Xbp1 branch of the UPR was not unequivocally detected in *Rp^+/−^* mutants. *Rp^+/−^* cells may be sensitized to activate PERK by Xrp1-dependent changes in transcription of Perk, BiP, and other UPR genes (Figure 10).

If eIF2α phosphorylation level, or its effects on translation, are involved in human ribosomopathies, its manipulation might be therapeutic. It is notable that knock-out of CReP, one of the two mouse PPP1R15 homologs, causes anemia, similar to DBA (Harding et al., 2009; Da Costa et al., 2018), and that PERK-dependent eIF2α phosphorylation occurs in RpL22-deficient mouse αβ T-cells, and activates p53 there (Solanki et al., 2016). Thus, inhibitors of eIF2α phosphorylation could be explored as potential DBA drugs. TAF1B depletion, which also acted through Xrp1 and eIF2α phosphorylation in *Drosophila*, is a model of Treacher Collins Syndrome(Trainor, Dixon, & Dixon, 2009), and failure to release eIF6, leading to defective LSU maturation and 80S ribosome formation, causes Schwachman Diamond syndrome(Warren, 2018), two other ribosomopathies where potential contributions of eIF2α phosphorylation remain to be investigated.

### Differences in translation can cause competition between cells but indirectly, through Xrp1

Because eIF2α phosphorylation alone was sufficient to target cells for competitive elimination, it seemed at first that eIF2α phosphorylation was the mechanism by which Xrp1 caused cell competition, which often correlates with differences in cellular translation levels(Nagata et al., 2019). Importantly, since another group concluded that eIF2α phosphorylation in *Rp^+/−^* cells did not lead to cell competition(Baumgartner et al., 2021), our different conclusion is independently corroborated by the observation that haploinsufficiency for the *eIF2*γ gene, which encodes another subunit of eIF2, initiates cell competition as efficiently as *Rp* haploinsufficiency does(Ji et al., 2021). We found here, however, that eIF2α phosphorylation did not cause cell competition directly, but because phosphorylation of eIF2α was itself sufficient to activate Xrp1 expression, and therefore cell competition through other Xrp1 targets(Figure 10). Elimination of *eIF2*γ haploinsufficient cells is also Xrp1-dependent, as expected if it is Xrp1 that is the key regulator of cell competition downstream of eIF2 activity(Ji et al., 2021). Knock-down of factors directly involved in the translation mechanism further distinguished cell competition from differential translation levels. Like eIF2α phosphorylation, these defects induced Xrp1 expression, which was required for the cell competition observed. Altogether, these results confirmed that reductions in global translation only trigger cell competition when Xrp1 is induced (Figure 10).

Through Xrp1, translation factor knockdown in turn also led to eIF2α phosphorylation. This was responsible for some of the dependence of global translation on translation factors ie translation was partially restored in cells depleted for translation factors when Xrp1 was depleted or eIF2α dephosphorylated. Surprisingly, eIF6 and eEF1α1 knockdown seemed not to reduce global translation at all, other than through Xrp1.

### Transcriptional regulation of cell competition

How does Xrp1 mark cells for competitive elimination, if not through eIF2α phosphorylation and reduced translation? Here we confirm that Xrp1 is a sequence-specific transcriptional activator. One suggestion has been that *Rp^+/−^* cells experience oxidative stress, and that an oxidative stress response predisposes them to elimination by competition with wild type cells(Kucinski et al., 2017; Baumgartner et al., 2021; Recasens-Alvarez et al., 2021). Because our studies showed that GstD1-GFP, the oxidative stress reporter in previous studies, was probably activated directly by Xrp1-binding in *Rp^+/−^* cells, and that a Xrp1-site mutated reporter was inactive in *Rp^+/−^* cells although retaining the Nrf2-dependent ARE site, it is questionable whether *Rp^+/−^* cells in fact experience significant oxidative stress or Nrf2 activity. Instead, direct transcriptional targets of Xrp1 may predispose *Rp^+/−^* and other cells to elimination by competition with wild type cells (Figure 10).

### Xrp1 as a central orchestrator of cell competition

Our results reveal the central importance of Xrp1 as the driver of cell competition (Figure 10). Far from being expressed specifically in *Rp* mutants, we now find that Xrp1 is induced by multiple challenges, not only to ribosome biogenesis, such as depletion of the polI cofactor TAF1B or LSU maturation factor eIF6, but to ribosome function, both at the levels of initiation or elongation, leading to cell competition and to Xrp1-dependent eIF2α phosphorylation. Whereas RpS12, which is crucial for Xrp1 induction in *Rp^+/−^* cells, was also important for TAF1B-depleted cells, it was less important for cells where ribosome function was affected, suggesting distinct mechanisms for Xrp1 induction (Figure 10).

It will be important now to determine whether yet other examples of cell competition involve Xrp1. For example, cells mutated for Helicase at 25E (Hel25E), a helicase that plays roles in mRNA splicing and in mRNA nuclear export, are lost in competition with wild type cells(Nagata et al., 2019). Although this has been attributed to lower translation in Hel25A cells, another explanation could be that Hel25A depletion induces Xrp1 expression. Cells with other defects affecting translation are also reportedly disadvantaged in mosaics, including mutations of an eIF5A-modifying enzyme(Patel, Costa-Mattioli, Schulze, & Bellen, 2009), and mutations of a pre-rRNA processing enzyme(Zielke, Vaharautio, Liu, & Taipale). It would not be surprising if other conditions that lead to eIF2α phosphorylation, such as ER stress, nutrient deprivation, or viral infection(Ron & Walter, 2007; Hetz, 2012), also activate Xrp1 and are thereby marked for elimination by more normal neighbors (Figure 10). It will be interesting to determine whether any of these conditions could interfere with surveillance and removal of aneuploid cells, given the potential importance for tumor surveillance(Ji et al., 2021). For example, it was already observed that nutrient deprivation interferes with competition of *Rp^+/−^* cells(Simpson, 1979), and therefore could interfere with the removal of aneuploid cells.

It will be important in future to evaluate Xrp1 expression and function in other examples of cell competition. Had we not evaluated Xrp1 expression and function in PPP1R15-depleted cells, we could have concluded that eIF2α phosphorylation was the likely downstream effector of competition in *Rp* mutant cells, rather than an example of a further upstream stress that induces Xrp1, which appears to be the common driver of cell competition for multiple genotypes (Figure 10).

## Materials and Methods

### Experimental Animals

Fly strains were generally maintained at 25°C on yeast cornmeal agar. Yeast-glucose medium was generally used for mosaic experiments (Sullivan et al., 2000). Sex of larvae dissected for most imaginal disc studies was not differentiated.

### Clonal Analysis

Genetic mosaics were generated using the FLP/FRT system using inducible heat shock FLP (hsFLP) transgenic strains. For making clones through mitotic recombination using inducible heat shock FLP (hsFLP), larvae of *Rp^+/−^* genotypes were subjected to 10-20 min heat shock at 37°C, 60 ± 12 hours after egg laying (AEL) and dissected 72 hr later. For making clones by excision of a FRT cassette, larvae were subjected to 10-30 min heat shock at 37°C (details in Suppl. Data Table 1), 36 ± 12 AEL for wild type background or 60 ± 12 hours AEL for *Rp^+/−^* background, and dissected 72 hr later.

### Drosophila stocks

Full genotypes for all the experiments are listed in Suppl. Data Table 1. The following genetic strains were used: UAS-PPP1R15 (BL76250), UAS-PERK-RNAi (v110278 and v16427), UAS-Gcn2-RNAi (v103976), TRE-dsRED (37), P[GAL4-Act5C(FRT.CD2).P]S, P[UAS-His-RFP]3 (isolated from BL51308), UAS-TAF1B-RNAi (BL61957 and v105783), UAS-PPP1R15-RNAi (v107545 and BL 33011), UAS-w-RNAi (BL33623), UAS-CG6272-RNAi (BL33652), Xbp1-EGFP (33), UAS-eIF4G-RNAi (v17002), UAS-eEF2-RNAi (v107268), UAS-eEF1α1-RNAi (v104502), UAS-eIF5Α-RNAi (v101513), UAS-eIF6-RNAi (v108094). Other stocks are described in (Lee et al., 2018).

### Immunohistochemistry and Antibody Labeling

For most antibody labeling, imaginal discs were dissected from late 3rd instar larvae in 1xPBS buffer and fixed in 4% formaldehyde in 1x PEM buffer (1xPEM:100mM Pipes, 1mM EGTA, 1mM MgCl2, pH 6.9). For p-eIF2α and p-RpS6 detection, larvae were dissected in Drosophila S2 medium one by one and transferred immediately to fixative. Fixed imaginal discs were 3x washed in PT (0.2% Triton X-100, 1xPBS) and blocked for 1 hour in PBT buffer (0.2% Triton X-100, 0.5% BSA, 1x PBS). Discs were incubated in primary antibody in PBT overnight at 4°C, washed 3 times with PT for 5-10 min each and incubated in secondary antibody in PBT for 3-4 hours at room temperature, and washed 3 times with PT for 5-10 min. After washes, samples were rinsed in 1x PBS and the samples were incubated with the NuclearMask reagent (Thermofisher, H10325) for 10-15 min at room temperature. After washing 2x with 1x PBS the imaginal discs were mounted in VECTASHIELD antifade mounting medium (Vector Laboratories, H-1000). In experiments that we wanted to parallel process control samples on the same tube (e.g. Figure 5C vs 5J), we used male parents that had the genotypes hsFLP; TRE-RFP/(PPP1R15 or Xrp1RNAi or PERKRNAi); act>>Gal4, UAS-GFP and cross them with females from the RNAi of interest. The genotypes in the same tube were discriminated using RFP before the addition of the secondary antibody. We used the following antibodies for staining: rabbit anti-phospho-RpS6 at 1:200 (kindly given by A. Teleman, DKFZ) (1:200), rabbit-p62 (kindly provided by Dr Juhász Gábor), rabbit anti-phospho-eIF2α at 1:100 (Thermofisher, 44-728G), rabbit anti-Xrp1 at 1:200 (kindly provided by D. Rio), mouse anti-b-Galactosidase (J1e7, DSHB), rabbit anti-GFP, rabbit anti-active-Dcp1 (Cell Signaling Techonology Cat#9578, 1:100), Y10b(1:100) (Thermofisher, MA1-13017), RpS9 (1:100) (Abcam, ab117861), RpL9(1:100) (Abcam, ab50384), rabbit-anti-Rack1 (1:100) (Cell Signalling, D59D5), rabbit anti-hRpL10Ab (1:100) (Sigma, Cat# SAB1101199). Secondary Antibodies were Cy2- and Cy5-conjugates (Jackson Immunoresearch) and Goat anti-Rabbit Alexa Fluor 555 (A21429, Thermofisher). Previous experiments established that significant results could be obtained from 5 replicates, although many more were imaged in most cases. No calculations regarding sample sizes were performed. No outliers or divergent results were excluded from analysis.

### Image Acquisition and Processing

Confocal laser scanning images were acquired with a Leica Laser scanning microscope SP8 using 20x and 40x objectives. Images were processed using Image J1.44j and Adobe Photoshop CS5 Extended. Thoracic bristle images were recorded using Leica M205 FA and Leica Application Suite X.

### Measurement of in vivo translation

Translation was detected by the Click-iT Plus OPP Alexa Fluor® 594 or 488 Protein Synthesis Assay Kit (Thermofisher, C10457) as described earlier (Lee et al, 2018). Larvae were inverted in Schneider’s Drosophila medium (containing 10% heat inactivated Fetal Bovine Serum, Gibco) and transferred in fresh medium containing 1:1000 (20uM) of Click-iT OPP reagent. Samples were incubated at room temperature for 15 minutes and rinsed once with PBS. The samples were fixed in 4% formaldehyde in 1x PEM buffer (100mM Pipes, 1mM EGTA, 1mM MgCl2) for 20 min, washed once with 1x PBS and subsequently washed with 0.5% Triton in 1x PBS for 10 min and then incubated for 10 min with 3% BSA in 1x PBS. The Click reaction took place in the dark at room temperature for 30 min. Samples were washed once with the rinse buffer of the Click reaction kit, 2 minutes with 3% BSA in 1x PBS, incubated for 1 hour at room temperature with PBT (1x PBS, 0.2% Triton, 0.5% BSA) and after that incubated overnight with the primary antibodies at 4oC. Samples were washed 3x with PT buffer (1x PBS, 0.2% Triton) and the secondary antibody was added for 2 hrs in room temperature. After 3x washes with PT and 1x with 1x PBS, the samples were incubated with the Nuclear Mask reagent (1:2000) of the Click-iT kit for 30 min. After washing 2x with 1x PBS the imaginal discs were mounted in Vectashield. Confocal laser scanning images were acquired with a Leica Laser scanning microscope SP8.

### Northern Analysis

RNA extraction, northern blotting procedures, and 18S, 5.8S, tubulin and actin probes were as described(Lee et al., 2018). Previous studies established that significant results could be obtained from 3 biological replicates. A biological replicate represents an independent RNA isolation, gel, and blot experiment.

The following primers were used to amplify the new probes in this paper:

ITS2 probe:

5’-CTTTAATTAATTTTATAGTGCTGCTTGG-3’

5’-TAATACGACTCACTATAGGGTTGTATATAACTTTATCTTG-3’

28S probe:

5’-GCAGAGAGATATGGTAGATGGGC-3’

5’-TAATACGACTCACTATAGGGTTCCACAATTGGCTACGTAACT-3’

ITS1 probe

5’-GGAAGGATCATTATTGTATAATATC-3’

5’-TAATACGACTCACTATAGGGATGATTACCACACATTCG-3’

**7SL probe:**

5’-TCGACTGGAAGGTTGGCAGCTTCTG-3’

5’-TAATACGACTCACTATAGGGATTGTGGTCCAACCATATCG-3’

### Plasmid cloning

All the new plasmids described below were confirmed by DNA sequencing.

#### Control *Renilla* luciferase plasmid

The pGL3-Promoter Vector (Promega) was modified by replacement of the SV40 promoter by the *Drosophila* actin promoter from the pAct5.1/V5-His C vector (Thermo Scientific), and the firefly luciferase coding sequence by the Renilla luciferase (RLuc) coding sequence from the pIS1 plasmid (Addgene), yielding the pGL3-Rluc plasmid.

#### *Firefly* luciferase plasmids

The SV40 core promoter of the pGL3-Promoter Vector was by hsp70 and Xrp1core promoters, amplified from the pUAST vector (*Drosophila* Genomics Resource Center) and from wild-type *Drosophila* genomic DNA respectively, using primers with XhoI and HindIII restriction sites. The resulting pGL3-H and pGL3-X plasmids were digested with Xho1 for insertion of annealed complementary oligonucleotides containing multiple copies of Target 1, Target 2, Target 3, or shuffled Target 1 or Target 2 sequences, resulting in the p-GL3-H-T1, p-GL3-H-T2, p-GL3-H-T3, p-GL3-H-T1S, p-GL3-H-T2S, pGL3-X-T1, pGL3-X-T2, pGL3-X-T3, pGL3-X-T1S, and pGL3-X-T2S plasmids.

#### Inducible expression plasmids

The Xrp1 (with and without its 3’UTR sequence) and Irbp18 (CG6272) coding regions were amplified from pUAST-Xrp1-HA and pUAST-CG6272 (Blanco et al., 2020), and inserted into pMT/V5-His A (Thermo Scientific) using XhoI and SpeI target sites, resulting in 3 inducible protein plasmids: pMT-Xrp1^HA^Δ3’UTR, pMT-Xrp1^HA^ and pMT-Irbp18^V5/His^. pMT-Xrp1^HA^ was not used further as it did not express Xrp1 protein in S2 cells.

### S2 cell culture and luciferase assays

*Drosophila* S2 cells from the *Drosophila* Genomics Resource Center (DGRC - stock#6) were cultured in Schneider’s medium (Thermo Scientific) supplemented with 10% Heat-Inactivated Fetal Bovine Serum (Thermo Scientific) at 25°C following the *General procedures for maintenance of Drosophila cell lines* from the DGRC. For luciferase assays, S2 cells were plated in 24-well plates, 5 x 10^5^ cells per well. After 24h cells were transfected with the indicated combination of control Rluc (0.15 ng/well), protein expression (15 ng/well) and target (4.5 ng/well) plasmids using TransIT-2020 Transfection Reagent (Mirus) following the manufacturer’s instructions. After 24h copper sulfate was added to a final concentration of 0.35 mM. After a further 24h cells were lysed and *Renilla* and *Firefly* luciferases’ activity measured with a luminometer, following the instructions from the Dual-Luciferase Reporter Assay System (Promega). *Firefly* signal was normalized to the internal *Renilla* control. Each transfection was performed in triplicate, and experiments performed independently at least 3 times.

## ACKNOWLEDGEMENTS

We thank Don Rio, Juhász Gábor and Aurelio Teleman for antibodies and Dirk Bohman, Katerina Papanikolopoulou, Hyung Don Ryoo, Efthimios Skoulakis and Eleni Tsakiri for other reagents. We thank Christos Delidakis, Nikolaos Konstantinides, Amit Kumar, Sudershana Nair, Venkateswara Reddy, Efthimios Skoulakis, and Deepika Vasudevan for comments on an earlier version of the manuscript. We thank Andreas Stasinopoulos for discussions, and Hyung Don Ryoo for sharing unpublished results. This work was supported by NIH grant GM120451 to NEB. Drosophila stocks were obtained from the Bloomington Drosophila Stock Center and Vienna Stock Resource Center (supported by NIH P40OD018537). Confocal microscopy was performed in the Analytical Imaging Facility of the Albert Einstein College of Medicine (supported by the NCI P30CA013330) using the Leica SP8 microscope acquired through NIH SIG 1S10 OD023591. Some data in this paper are from a thesis submitted in partial fulfillment of the requirements for the Degree of Doctor of Philosophy in the Biomedical Sciences, Albert Einstein College of Medicine.

---

**<u>Figure 1 source data 1</u>**

Full and unedited blots corresponding to panel A.

**<u>Figure 1 source data 2</u>**

Northern data underlying panels B-E.

**Figure 1 Supplement 1.**
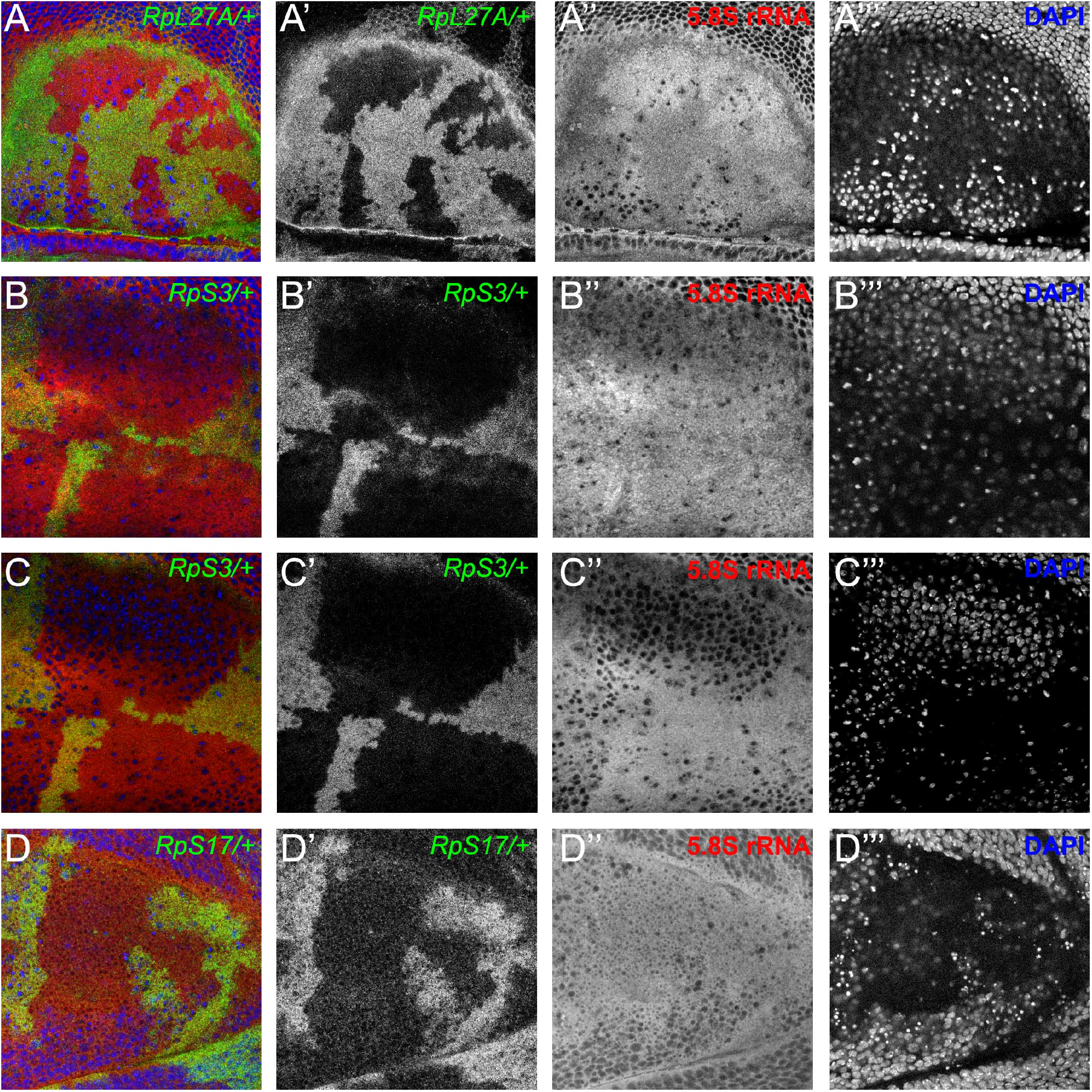
Cytoplasmic location of ribosome components. Specimens including those from Figure 1F,G showing the nuclear channel that was recorded simultaneously in many experiments. A) This very apical confocal plane passes through only a few nuclei, particularly in wild type cells, verifying that the anti-5.8S rRNA signal is predominantly cytoplasmic, consistent with mature LSU. B) This very apical confocal plane largely excludes nuclei, verifying that the anti-5.8S rRNA signal is predominantly cytoplasmic, consistent with mature LSU. C) This slightly less apical focal plane predominantly grazes nuclei only of wild type cells, verifying that little anti-5.8S rRNA signal is nuclear. D) This very basal confocal plane, which largely excludes nuclei for wild type regions, shows no discernible difference in cytoplasmic anti-5.8S rRNA signal between wild type and *RpS17*/+ cells.

**Figure 1 Supplement 2.**
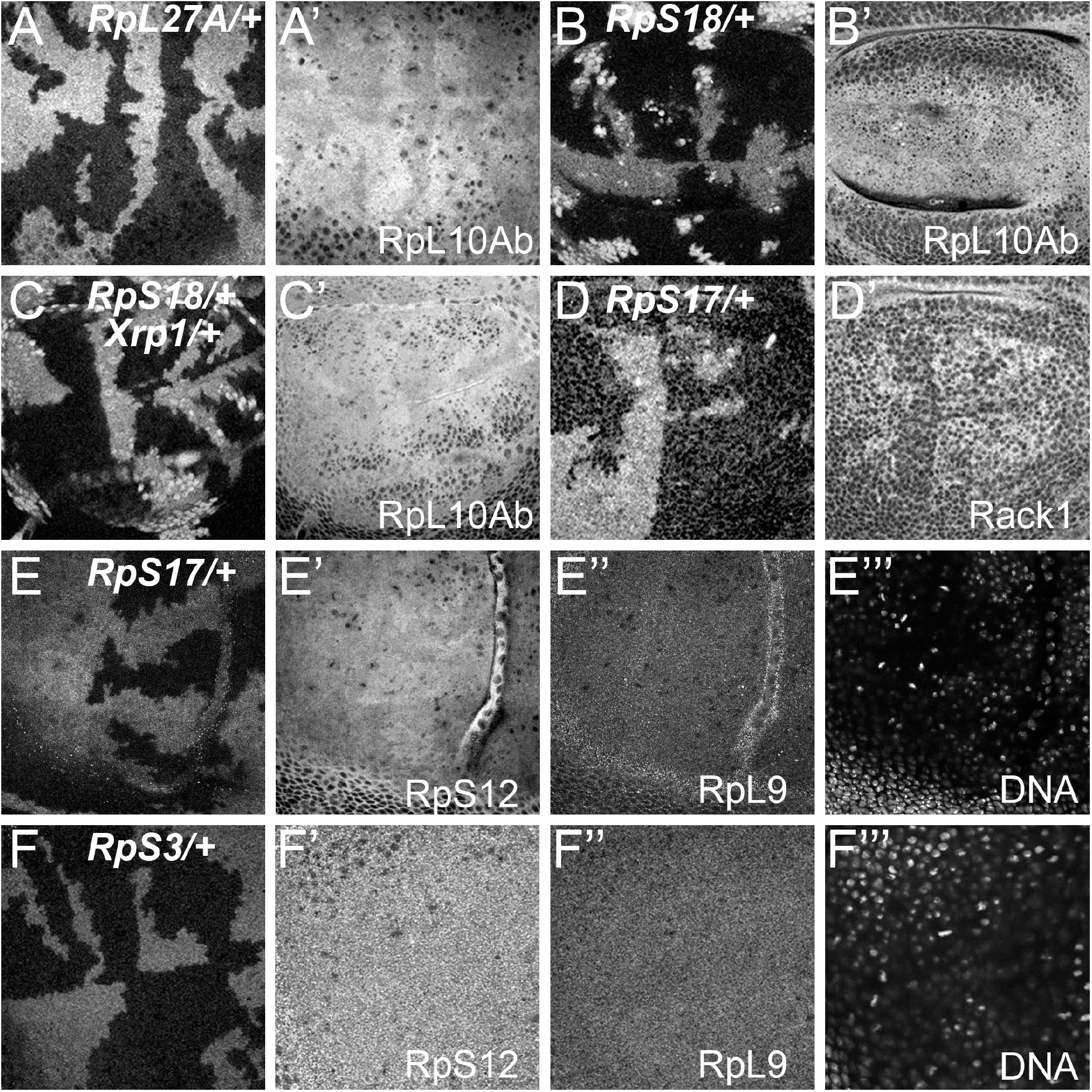
More examples of ribosome levels in *Rp* mutant genotypes. Panels A-F show mosaic wing discs containing unlabelled *Rp^+/+^* cells and labelled cells of indicated *Rp^+/−^* genotypes. Panels A’-F’, E’’ and F’’ show these same discs labelled with the antibodies against the indicated Rp. Panels E’’’ and F’’’ indicate DNA, in these cases indicating an apical confocal plane above most nuclei. A) *RpL27A^+/−^*cells contained less of the LSU component RpL10Ab. B) *RpS18^+/−^*cells contained more of the LSU component RpL10Ab. C) *RpS18^+/−^Xrp1^+/−^* cells contained more of the LSU component RpL10Ab than *RpS18^+/+^Xrp1^+/−^* cells. D) *RpS17^+/−^*cells contained less of the SSU component Rack1. E) *RpS17^+/−^*cells contained less of the SSU component RpS12 but levels of the LSU component RpL9 were indistinguishable from *RpS17^+/+^*cells. F) *RpS3^+/−^*cells contained levels of RpS12 and RpL9 indistinguishable from *RpS3^+/+^*cells.

**<u>Figure 2 source data 1</u>**

Full and unedited blots corresponding to panel B

**<u>Figure 2 source data 2</u>**

Full and unedited blots corresponding to panel C

**<u>Figure 2 source data 3</u>**

Full and unedited blots corresponding to panel D

**Figure 2 Supplement 1.**
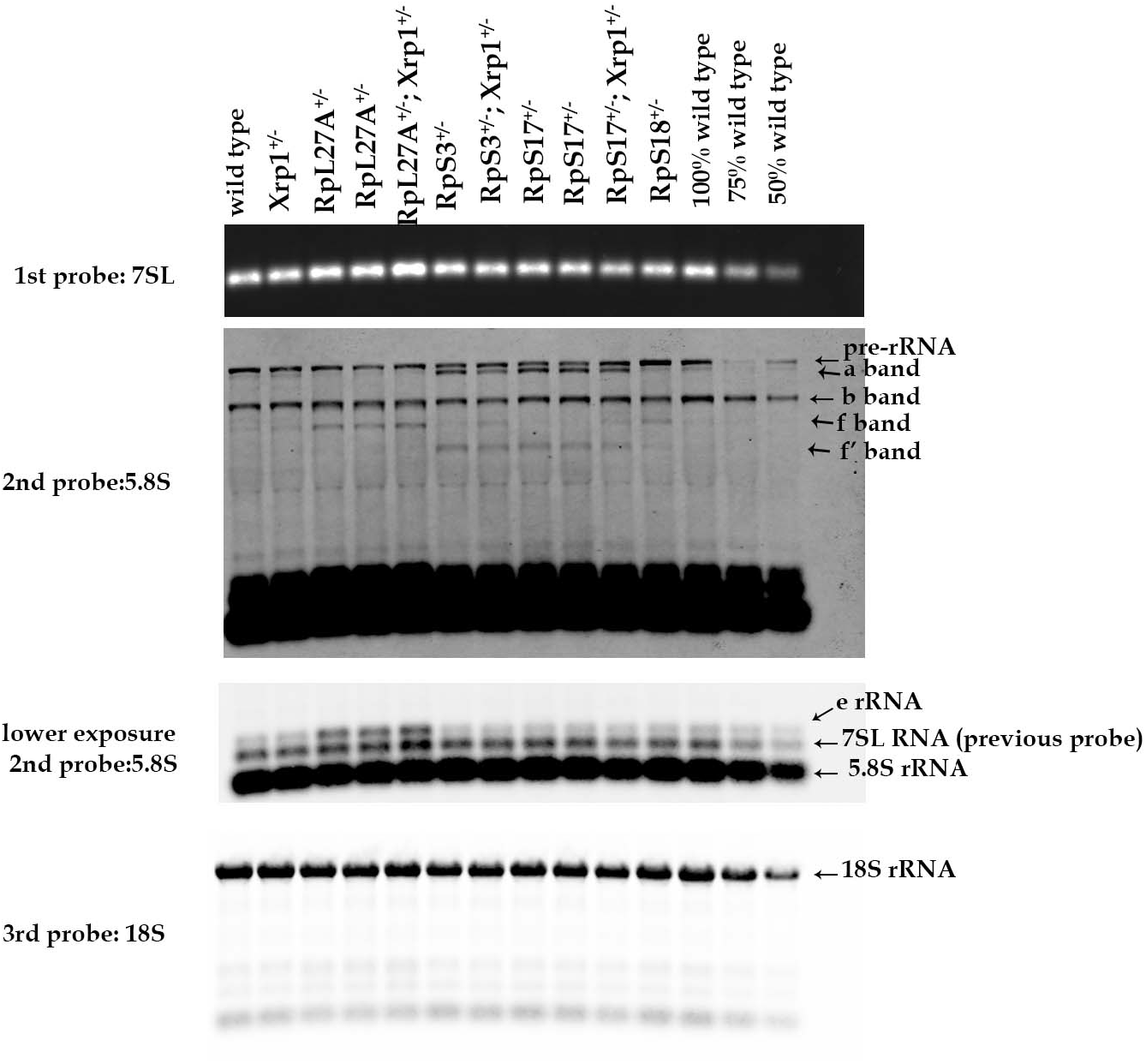
Additional northern blots detecting rRNA intermediates. Northern blots of total RNA purified from wild-type and Rp+/− wing discs, reprobed with 5.8S probe and then 18S probe after an initial 7SL probe.

**<u>Figure 2 Supplement 1 Source data 1</u>**

Full and unedited blots corresponding to Figure 2 supplement 1

**Figure 2 Supplement 2.**
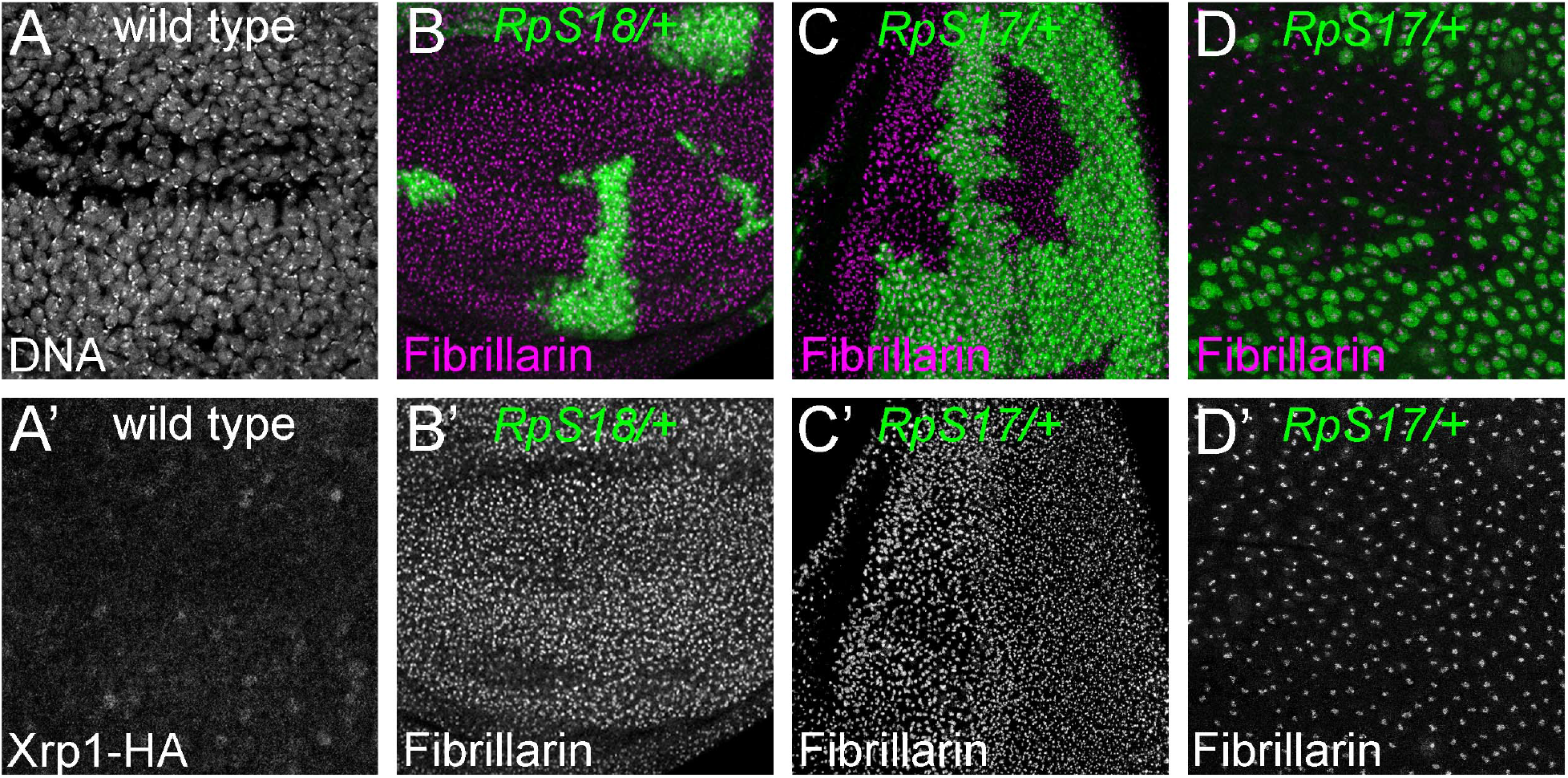
Nucleoli in wild type and *Rp* mutant cells. A) Confocal section of Xrp1-HA wing disc showing many nuclei labelled for DNA. A’) HA labeling reveals minimal Xrp1 expression in this otherwise wild type wing disc. Nucleoli are not obviously labelled. B) Mosaic wing disc containing *RpS18*^+/−^ cells (green). Anti-fibrillarin labeling of nucleoli reveals no obvious differences between *RpS18*^+/−^ and *RpS18*^+/+^ cells (projected in magenta; see also B’). C) Mosaic eye disc containing *RpS17*^+/−^ cells (green). Anti-fibrillarin labeling of nucleoli reveals no obvious differences between *RpS17*^+/−^ and *RpS17*^+/+^ cells (projected in magenta; see also C’).D) Peripodial membrane from mosaic eye eye disc containing *RpS17*^+/−^ cells (green). Anti-fibrillarin labeling of nucleoli reveals no obvious differences between *RpS17*^+/−^ and *RpS17*^+/+^ cells (projected in magenta; see also D’).

**Figure 3 Supplement 1.**
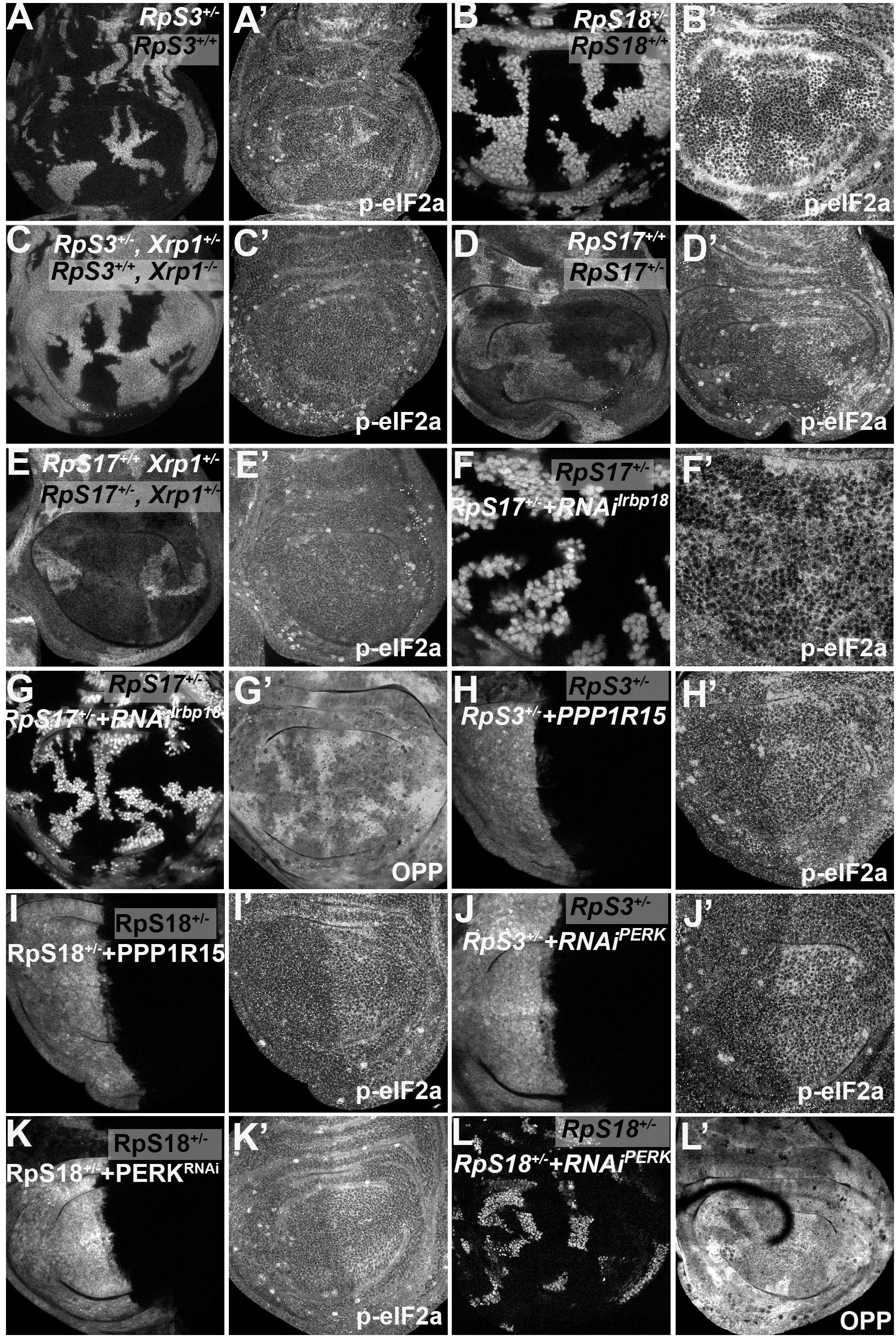
eIF2α phosphorylation in *Rp*^+/−^ cells depends on Xrp1 and Irbp18. Panels A-L show single confocal planes from third instar wing imaginal discs. A) Mosaic of *RpS3*^+/−^ and *RpS3*^+/+^ cells. p-eIF2α levels were increased in *RpS3*^+/−^ cells (see A’). B) Mosaic of *RpS18*^+/−^ and *RpS18*^+/+^ cells. p-eIF2α levels were increased in *RpS18*^+/−^ cells (see B’). C) Mosaic of *RpS3*^+/−^ *Xrp1^+/−^* and *RpS3*^+/+^ *Xrp^−/−^* cells. p-eIF2α levels were unaffected in *RpS3*^+/−^ cells when *Xrp1* was mutated(see C’). D) Mosaic of *RpS17*^+/−^ and *RpS17*^+/+^ cells (the latter labeled more brightly, having two copies of β-gal transgene). p-eIF2α levels were increased in *RpS17*^+/−^ cells (see D’). E) Mosaic of *RpS17*^+/−^ and *RpS17*^+/+^ cells in an *Xrp1^+/−^* wing disc, (*RpS17*^+/+^ labeled more brightly, having two copies of β-gal transgene).. p-eIF2α levels were unaffected in *RpS17*^+/−^ cells (see E’). F) Labelled clones of cells expressing Irbp18 RNAi in a *RpS17*^+/−^ wing disc. p-eIF2α levels were reduced by Irbp18 knock-down (see F’). G) Labelled clones of cells expressing Irbp18 RNAi in a *RpS17*^+/−^ wing disc. Translation rate was restored by Irbp18 knock-down (see G’). H) Labelled cells over-expressing PPP1R15 in the posterior compartment of a *RpS3*^+/−^ wing disc. p-eIF2α levels were reduced by PPP1R15 over-expression (see H’). I). Labelled cells over-expressing PPP1R15 in the posterior compartment of a *RpS18*^+/−^ wing disc. p-eIF2α levels were reduced by PPP1R15 over-expression (see I’). J) Labelled cells expressing PERK RNAi in the posterior compartment of a *RpS3*^+/−^ wing disc. p-eIF2α levels were reduced by PERK knock-down (see J’). K). Labelled cells expressing PERK RNAi in the posterior compartment of a *RpS18*^+/−^ wing disc. p-eIF2α levels were reduced by PERK knock-down (see K’). L) Labelled clones of cells expressing PERK RNAi in a *RpS18*^+/−^ wing disc. Translation rate was restored by PERK knock-down (see L’).

**Figure 4 Supplement 1.**
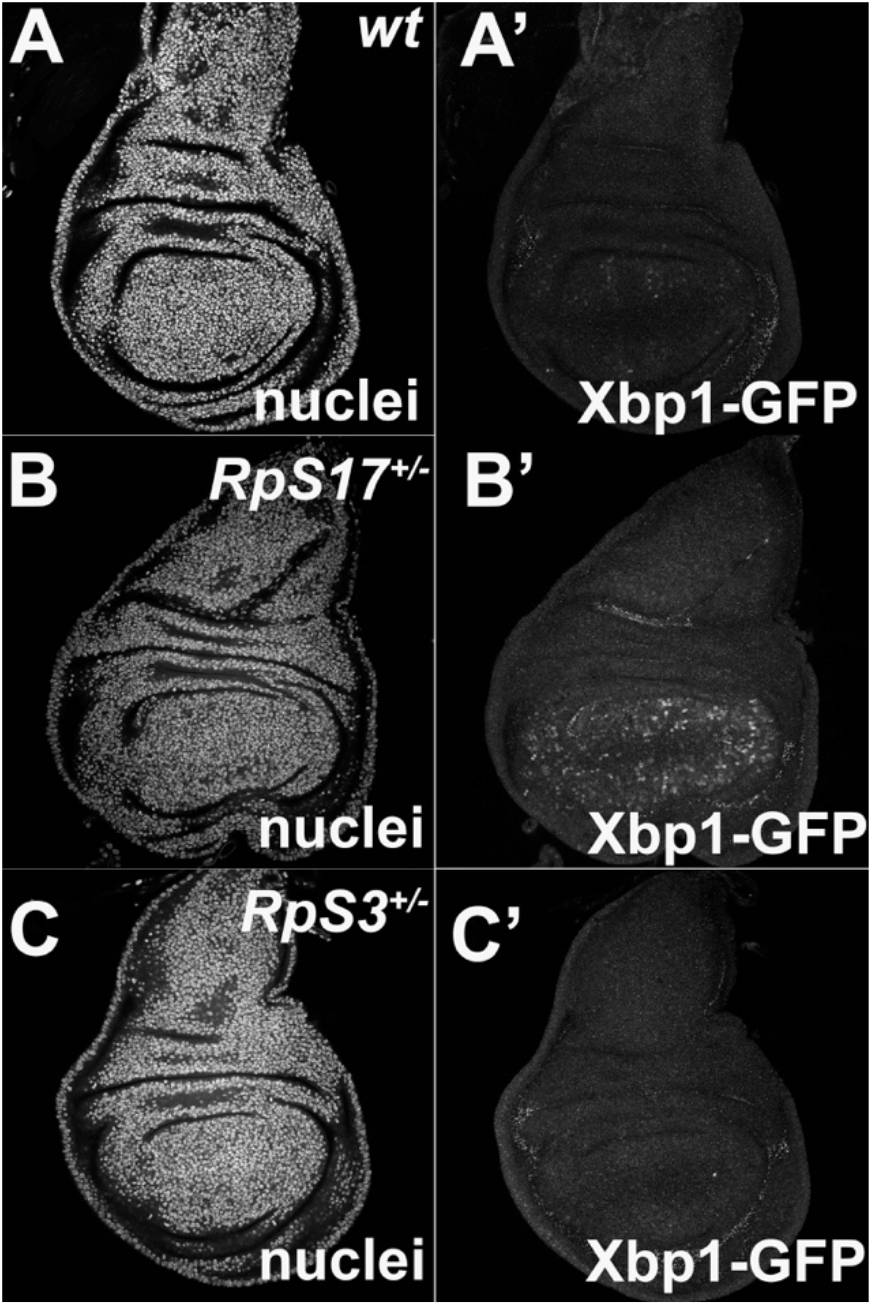
Little UPR detected in *Rp*^+/−^ wing discs. Panels show single confocal planes from third instar wing imaginal discs expressing the UPR reporter UAS-Xbp1-GFP in the wing pouch under *nub-Gal4* control. GFP expression indicates an unfolded protein response. A). Little evidence for UPR in wild type wing discs. B) Elevated UPR in *RpS17^+/−^* wing discs. C) Little evidence for UPR in *RpS3^+/−^* wing discs.

**Figure 5 Supplement 1.**
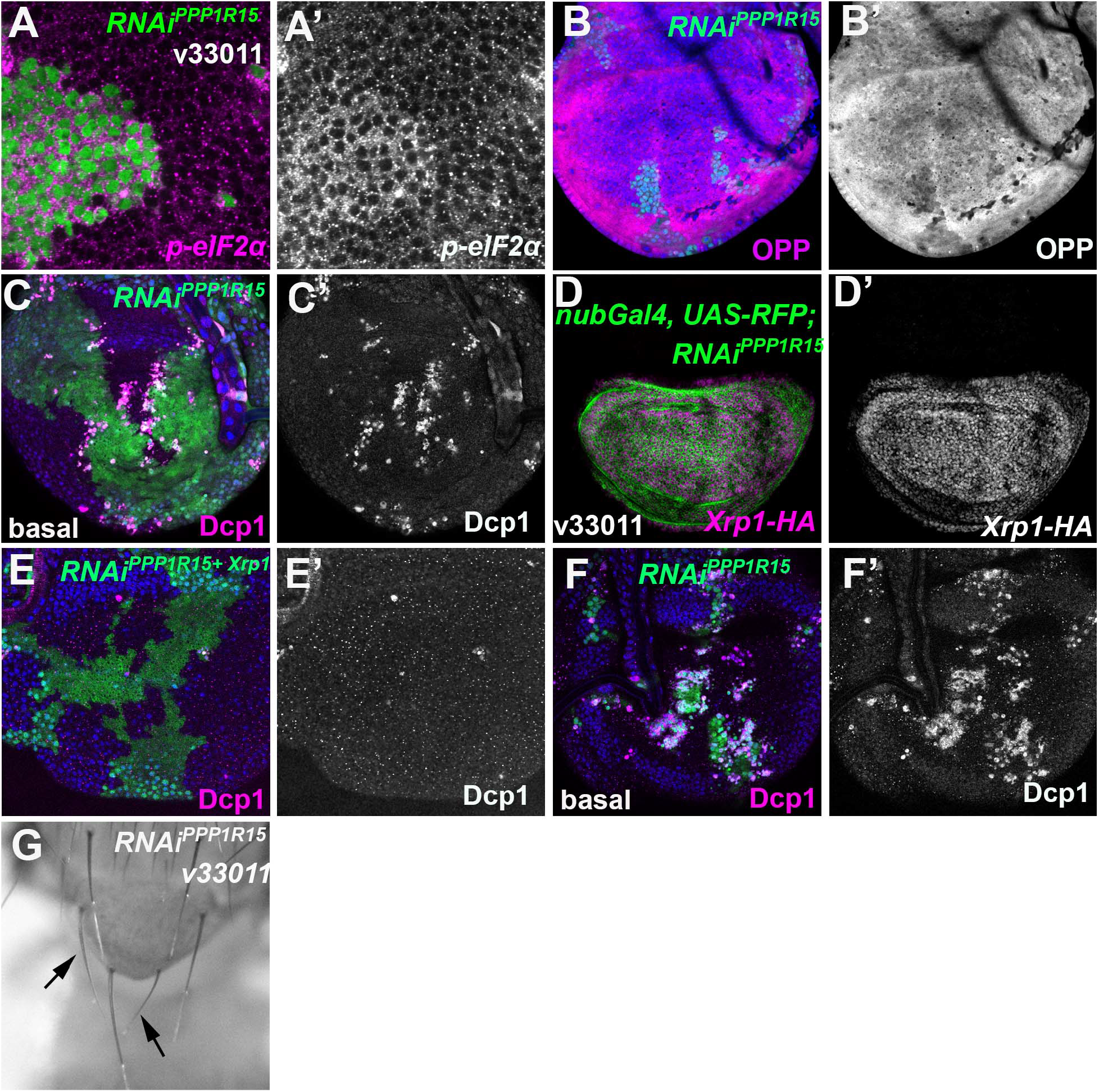
eIF2α phosphorylation induces Xrp1 expression, cell competition, and small bristles. Single confocal planes from third instar wing imaginal discs. A) Clones expressing PPP1R15-RNAi had increased p-eIF2α levels (A’). B) Clones expressing PPP1R15-RNAi had reduced translation (OPP) (B’). C) Basal section of the same disc shown in Figure 5E. More dying PPP1R15-RNAi cells labeled for active caspase (magenta, see also C’) accumulate basally at the boundaries with the wild type cells. D) *nub-Gal4* driving *PPP1R15* RNAi in the wing pouch (green) led to Xrp1-HA expression (magenta; see also D’). E) Basal confocal section of the wing disc also shown in Figure 5H, mosaic for cells expressing *PPP1R15* RNAi and Xrp1 RNAi (green). Even at these basal levels, apoptosis was almost completely rescued by Xrp1 knockdown (magenta, see also E’). F) Basal confocal section of wing disc mosaic for wild type cells and cells expressing *PPP1R15* RNAi also shown in Figure 5I, a parallel control to panel E. Note extensive cell death basally (magenta, see also F’), as well as smaller clone size (green). G) Minute-like short, thin bristles (arrows) on adults when PPP1R15 was depleted in clones contrast with the normal contralateral bristles.

**Figure 6 Supplement 1.**
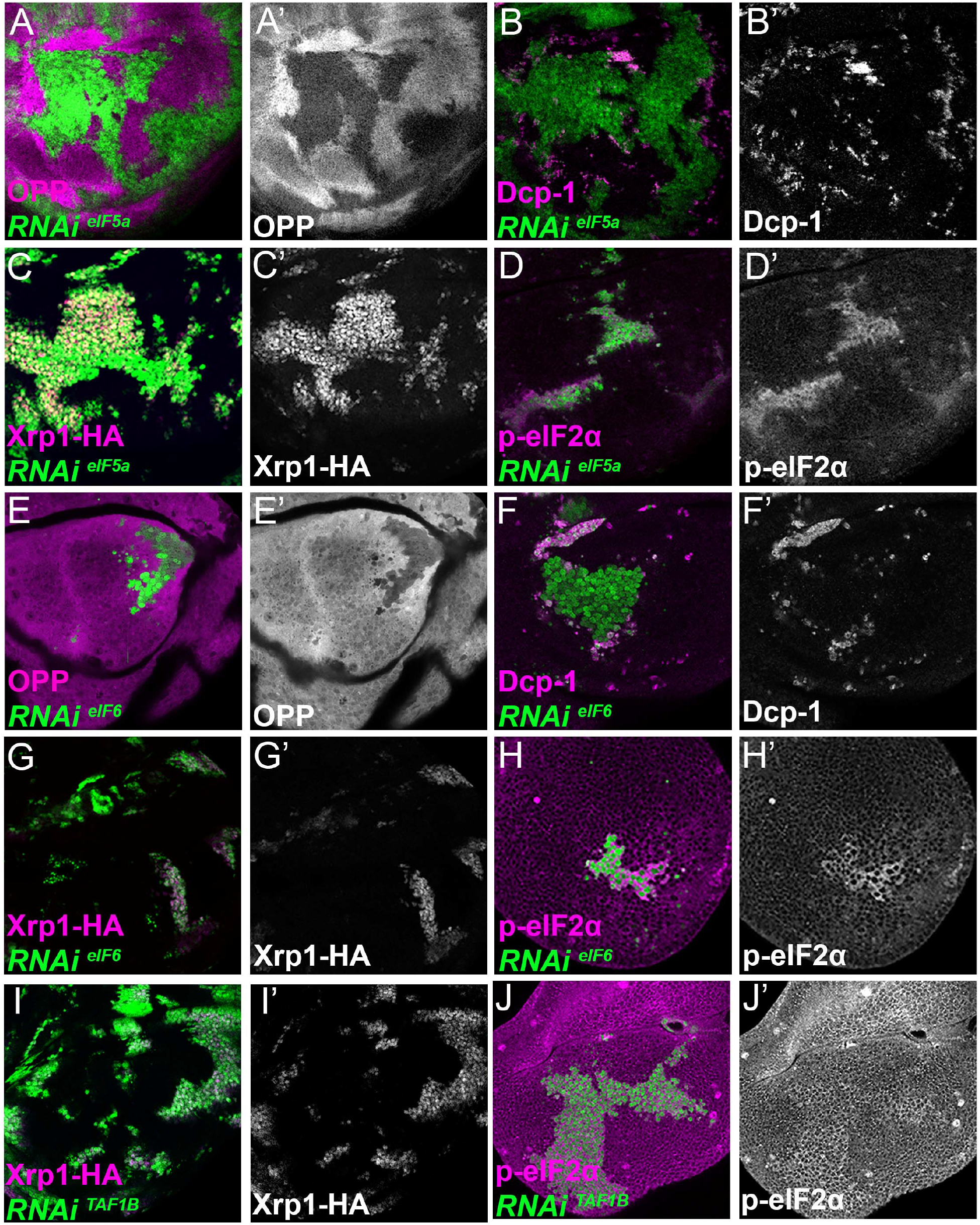
Xrp1 expression, eIF2α phosphorylation, reduced translation, and cell competition after depletion of additional translation factors. Clones of cells depleted for translation factors are shown in green. In each case, translation factor depletion reduced translation rate, resulted in competitive cell death at interfaces with wild type cells, induced Xrp1-HA expression, and led to eIF2α phosphorylation. Translation rate, dying cells (activated caspase Dcp1), Xrp1-HA and p-eIF2α are indicated in magenta and in separate channels as labelled. A-D) Clones expressing RNAi for eIF5A. E-H) Clones expressing RNAi for eIF6. I,J) Clones expressing RNAi for TAF1B.

**Figure 6 Supplement 2.**
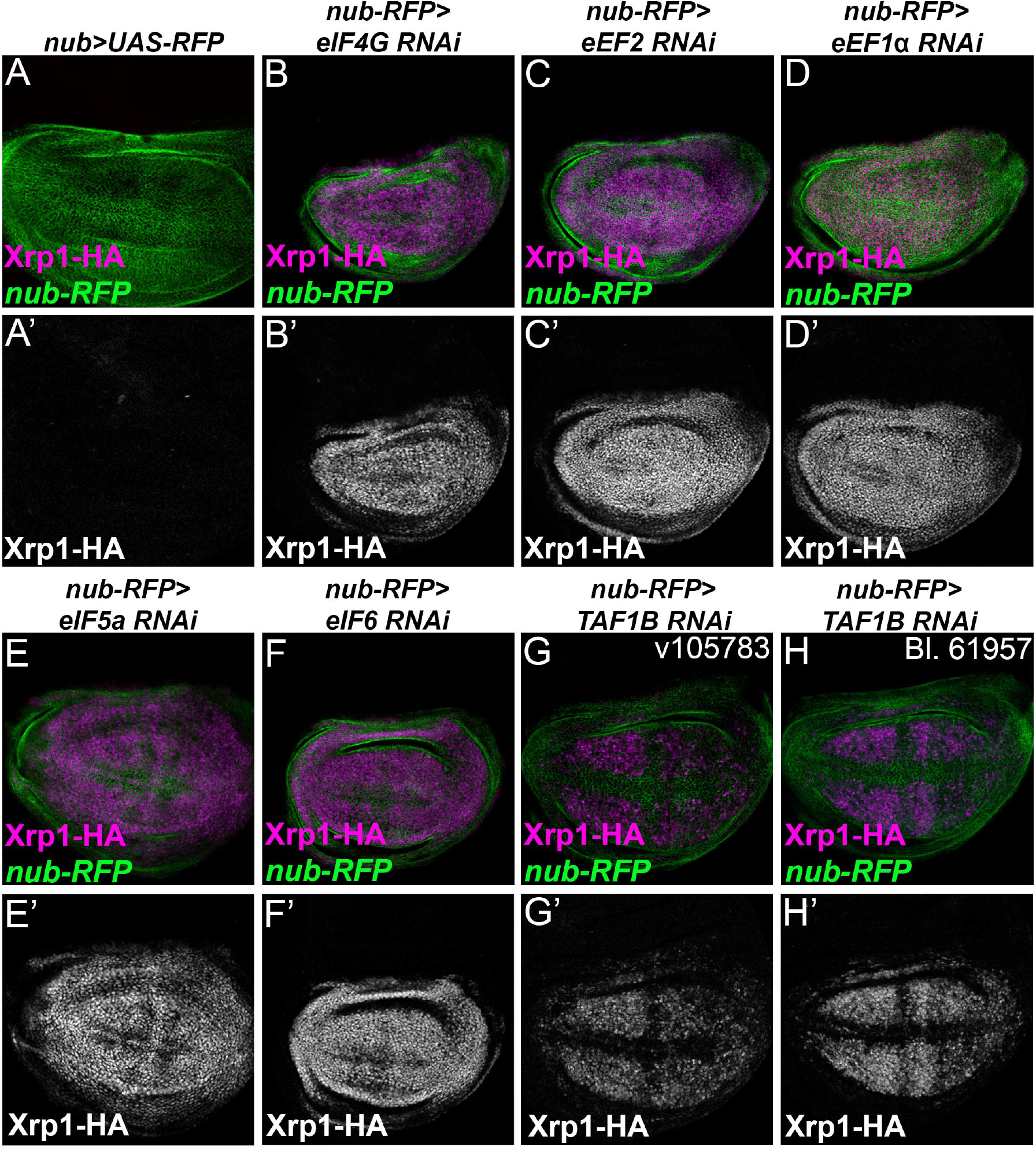
Translation factor knock-down induces Xrp1 expression. Knock down of translation factors using nub-Gal4 to drive RNAi in the wing pouch (green) induced Xrp1-HA expression (magenta, and separate channels as indicated). In the control of nub-Gal4 expressing RFP alone, negligible Xrp1-HA was detected (see A’). B) eIF4G-RNAi. C) eEF2-RNAi. D) eEF1α1-RNAi. E) eIF5A-RNAi. F) eIF6-RNAi. G) TAF1B-RNAi H) TAF1B-RNAi.

**Figure 7 Supplement 1.**
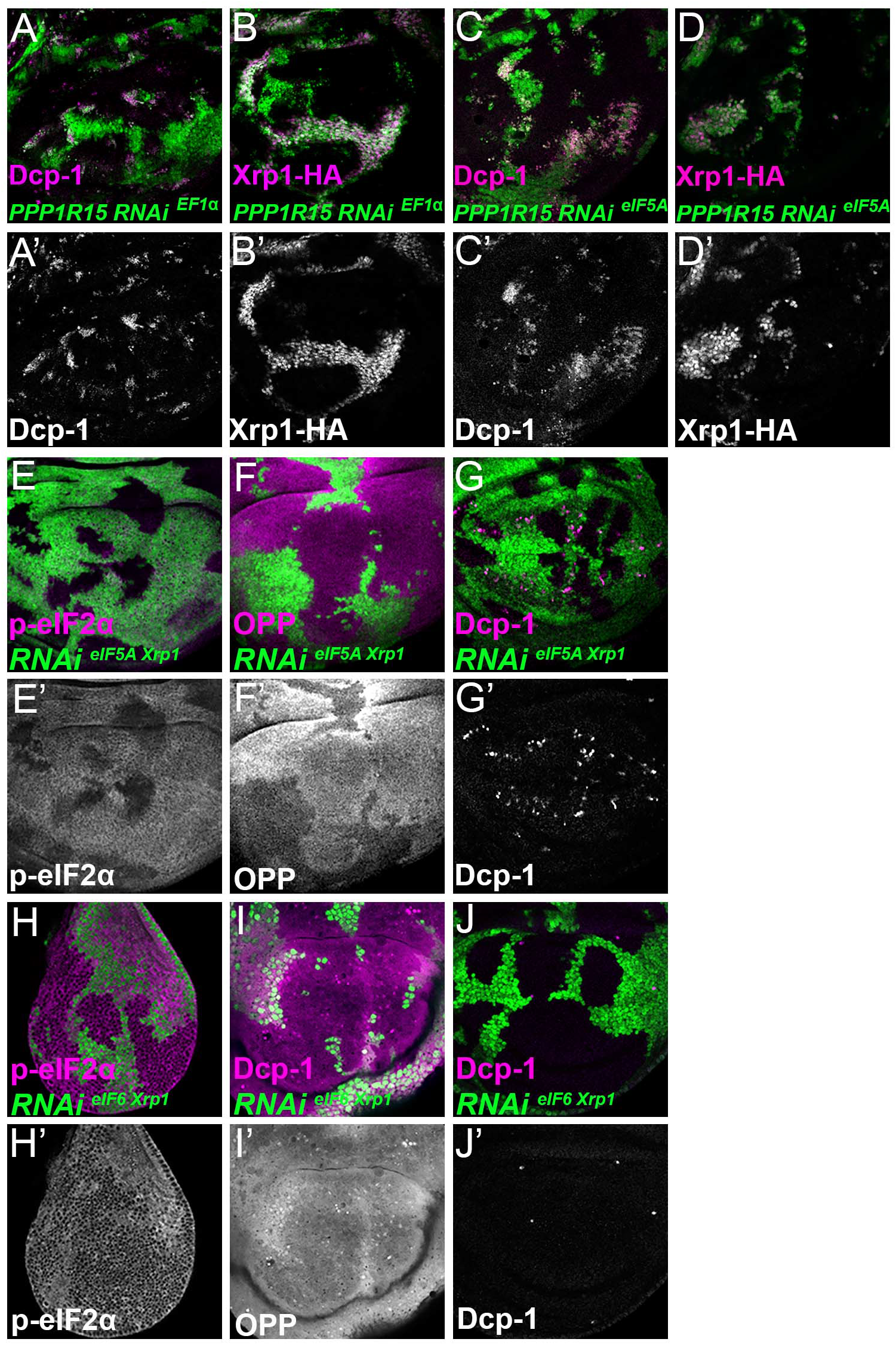
Single confocal planes from third instar wing imaginal discs. p-eIF2α levels, translation rate (ortho-propargyl puromycin), dying cells (activated caspase Dcp1) and Xrp1-HA are indicated in magenta and in separate channels as labelled. (A-D) Clones of cells depleted for translation factors which also overexpress PPP1R15 are shown in green. PPP1R15 overexpression did not reduce Xrp1-HA levels or rescue competitive cell death in the cells depleted for eEF1α (A,B) or eIF5A (C,D). (E-G) Clones of cells depleted eIF5A that also express RNAi for Xrp1 are shown in green. Xrp1-depletion did not reduce eIF2α phosphorylation or restore translation levels, but it reduced levels of competitive cell death (compare panel C and Figure 6B) (H-J) Clones of cells depleted eIF6 that also express RNAi for Xrp1 are shown in green. Xrp1 depletion in cells expressing eIF6 reduced eIF2α phosphorylation and rescued competitive cell death. Translation rates were restored at least to wild type levels.

**Figure 7 Supplement 2.**
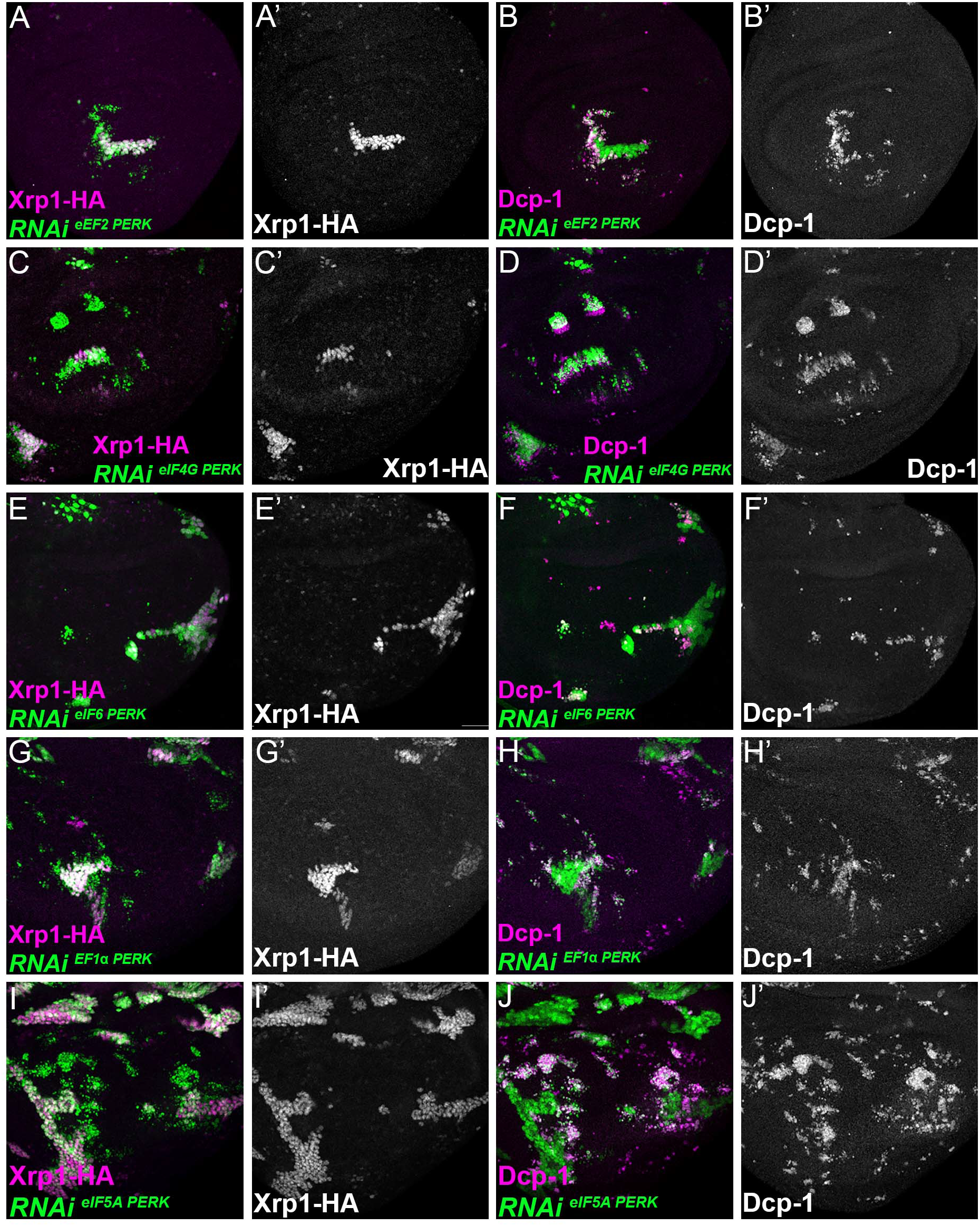
Single confocal planes from third instar wing imaginal discs. Dying cells (activated caspase Dcp1) and Xrp1-HA are indicated in magenta and in separate channels as labelled. Clones of cells depleted for translation factors which also overexpress PERK-RNAi are shown in green. In no case did PERK depletion affect Xrp1-HA induction or suppress competitive cell death. A,B) Clones expressing RNAi for both eEF2 and PERK. C,D) Clones expressing RNAi for both eIF4G and PERK. E,F) Clones expressing RNAi for both eIF6 and PERK. G,H) Clones expressing RNAi for both eEF1α1 and PERK. I,J). Clones expressing RNAi for both eIF5A and PERK.

**<u>Figure 9 source data file 1</u>**

Luciferase data relevant to panels C,D.

**<u>Figure 9 source data file 2</u>**

GFP data relevant to panel J

**<u>Figure 9 source data file 3</u>**

GFP data relevant to panel O

**Figure 9 Figure Supplement 1.**
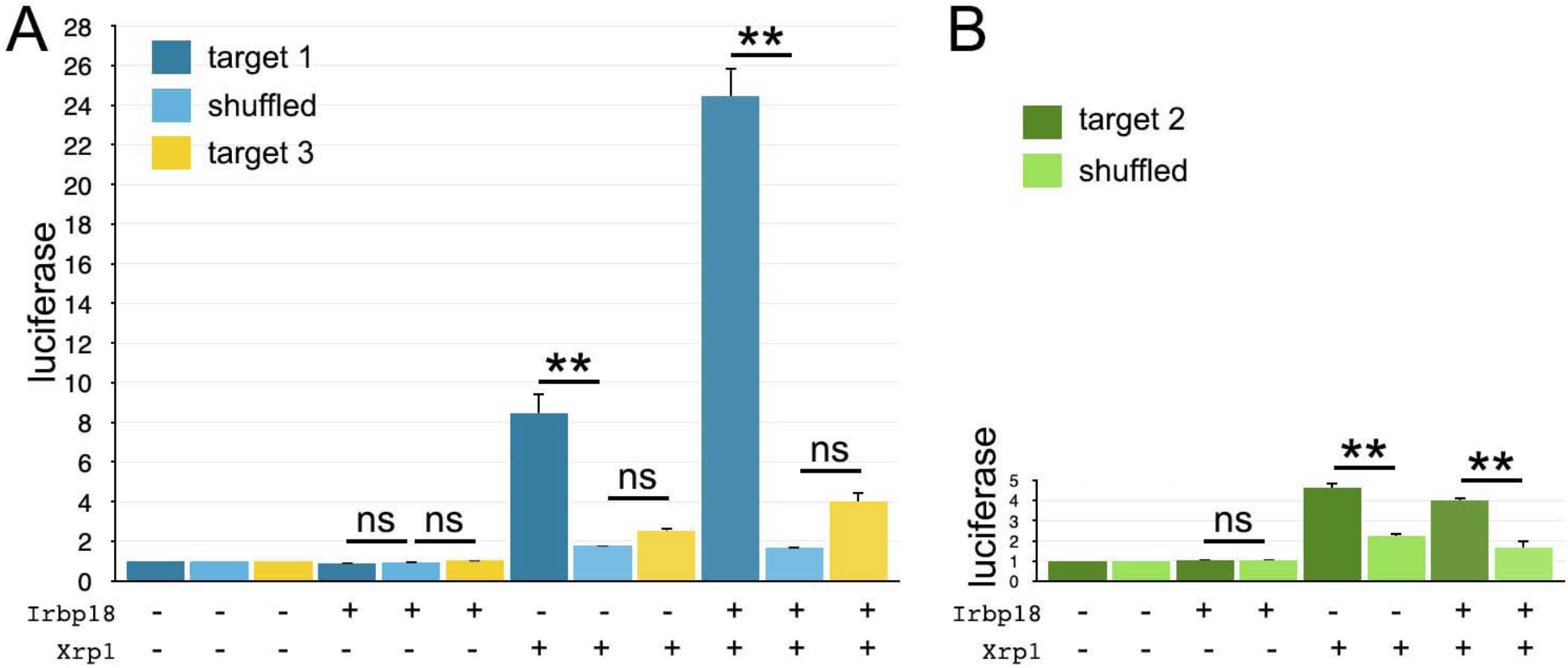
Luciferase assays with *hsp70*-based reporter plasmids. Fold change in the luciferase/renilla signal is shown for the remaining reporter constructs shown in Figure 9B. A) Target 1 conferred sequence-specific activation by transfected Xrp1 protein. Transfected Irbp18 alone had no effect, but synergized with Xrp1. Lesser activation by Target 3 was rendered not significant by correction for multiple testing, but the increase in both Xrp1 and Xrp1+Irbp18 samples suggests that it may nevertheless be real. B) Target 2 conferred sequence-specific activation by transfected Xrp1 protein. Transfected Irbp18 alone had no effect. It may be significant that Targets 1 and 3 are better matches to the in vivo-derived ChIP-Seq consensus than Target 2 (see Figure 9A). Statistics: 1-way ANOVA with Bonferroni-Holm correction for multiple testing was performed for the data shown in each of panels A,B. ns, p>0.05. **, p<0.01. Data were based on triplicate measurements from each of 3 biological replicates for each transfection. Exact p-values for comparisons between target 1 reporters and scrambled reporters (panel A) were: Padj=4.51, Padj=1.80×10^−6^, Padj=0 for Irbp18, Xrp1, and Irpb18+Xrp1 transfected cells respectively. Exact p-values for comparisons between target 2 reporters and scrambled reporters (panel A) were: Padj=2.81, Padj=4.22, Padj=0.225 for Irbp18, Xrp1, and Irpb18+Xrp1 transfected cells respectively. Exact p-values for comparisons between target 3 reporters and scrambled reporters (panel B) were: Padj=0.983, Padj=6.70×10^−6^, Padj=8.03×10^−6^, for Irbp18, Xrp1, and Irpb18+Xrp1 transfected cells respectively.

**<u>Figure 9 Figure Supplement 1 source data file 1</u>**

Luciferase measurements relevant to Figure 9 Supplement 1.

**Figure 9 Figure Supplement 2.**
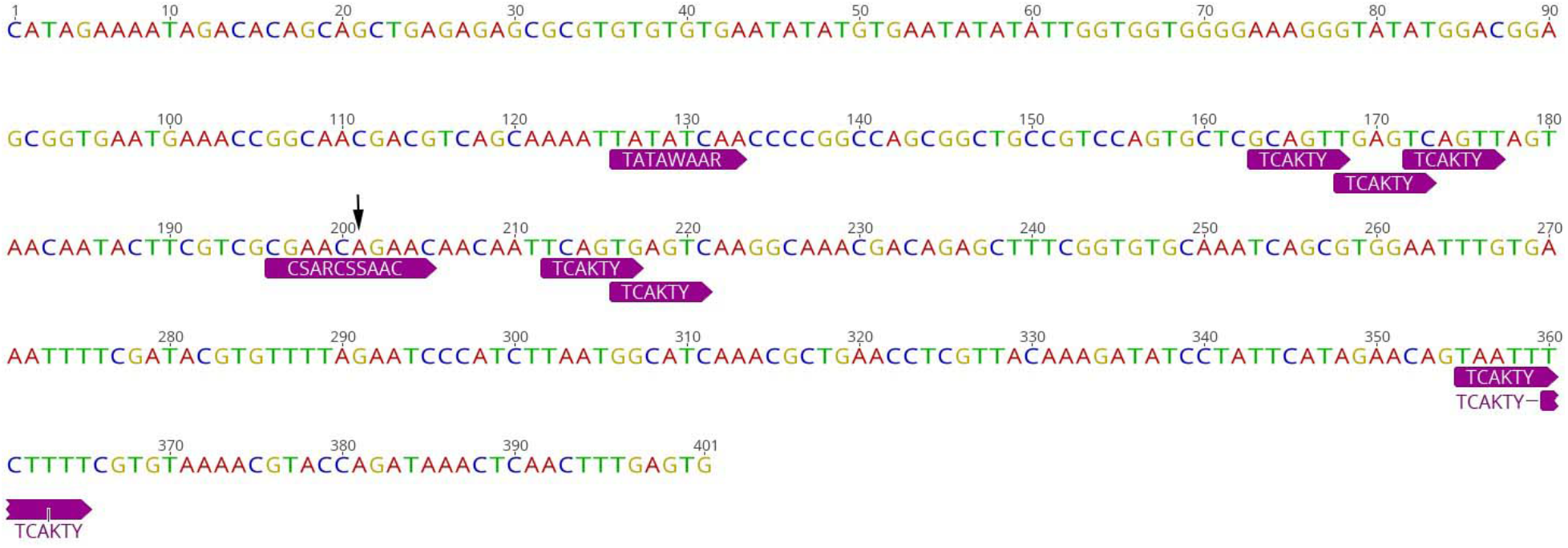
*Xrp1* promoter proximal sequences. The 400bp *Xrp1* core promoter sequence is shown. Transcription start site indicated by arrow. A variety of conserved promoter element sequences are indicated (Ohler, Liao, Niemann, & Rubin, 2002; Juven-Gershon & Kadonaga, 2010).

**Supplementary Table 1:**
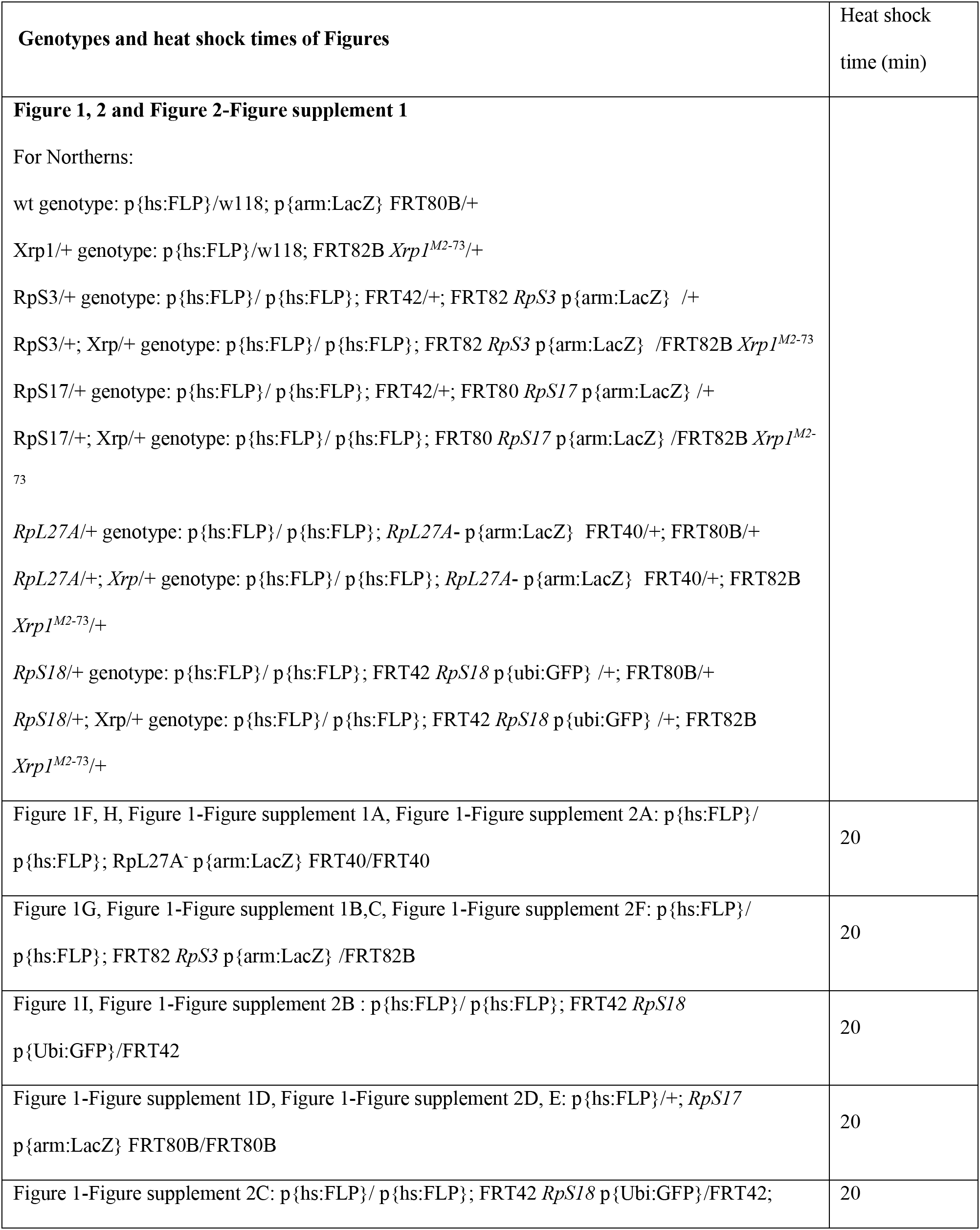
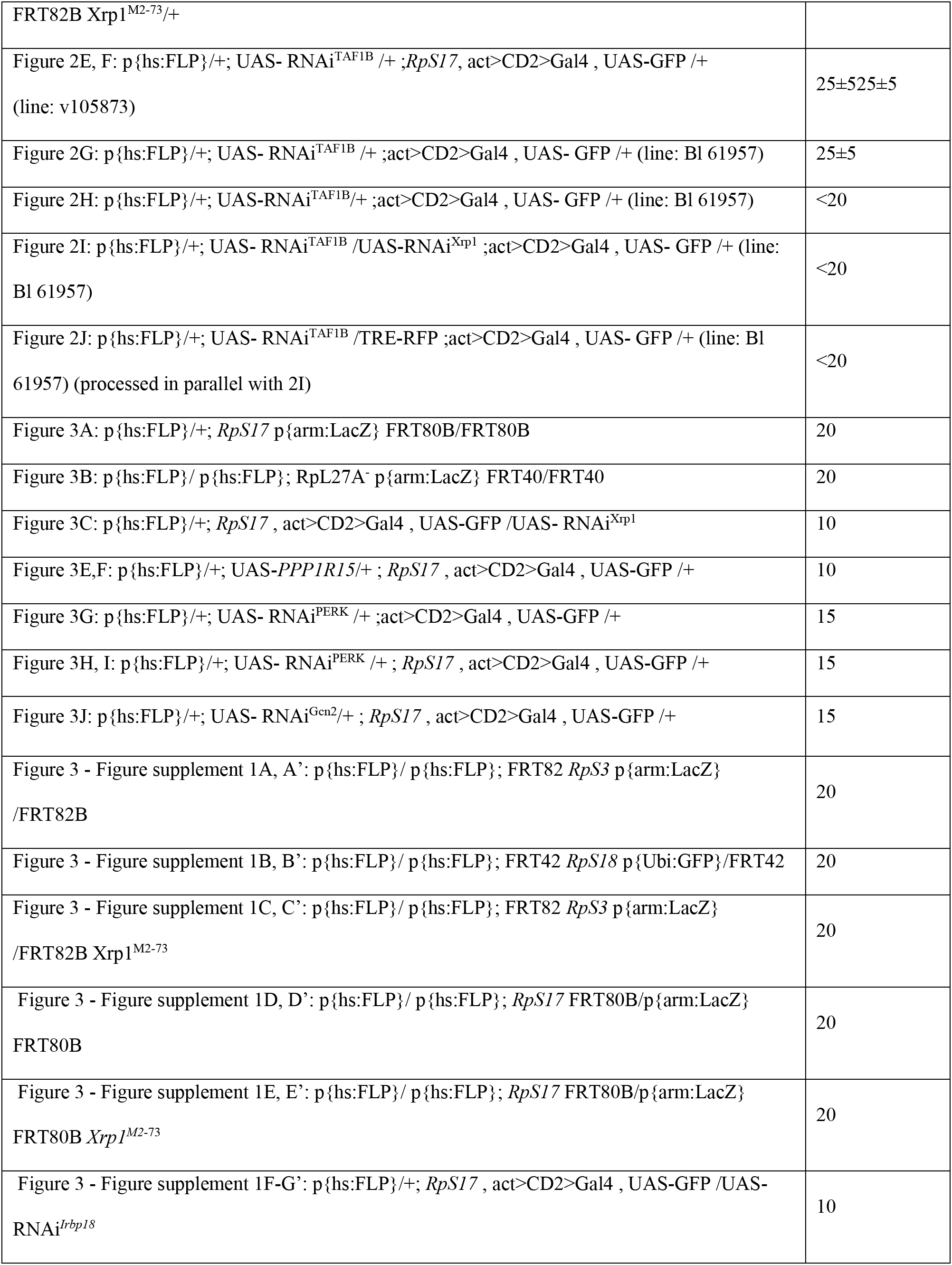
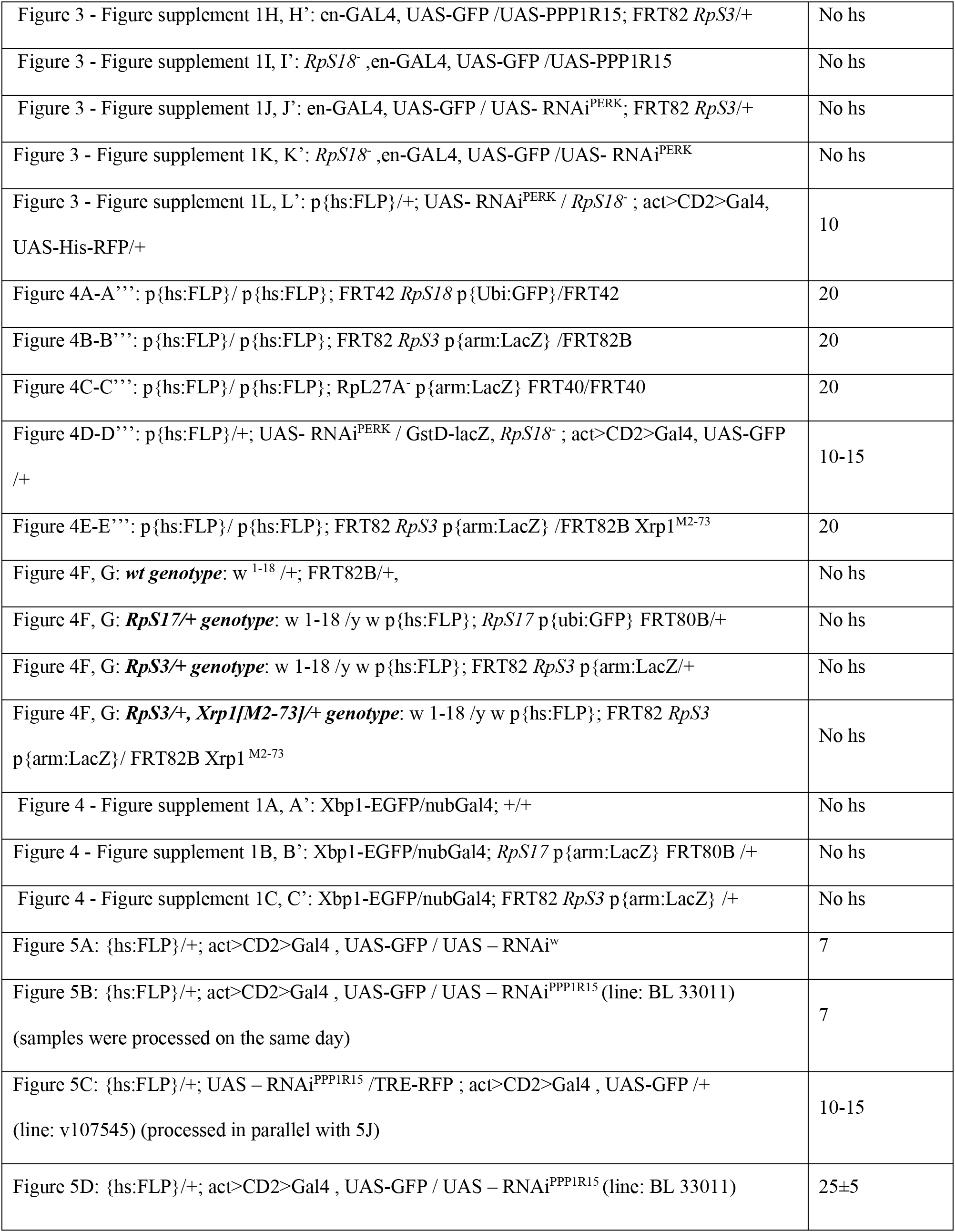
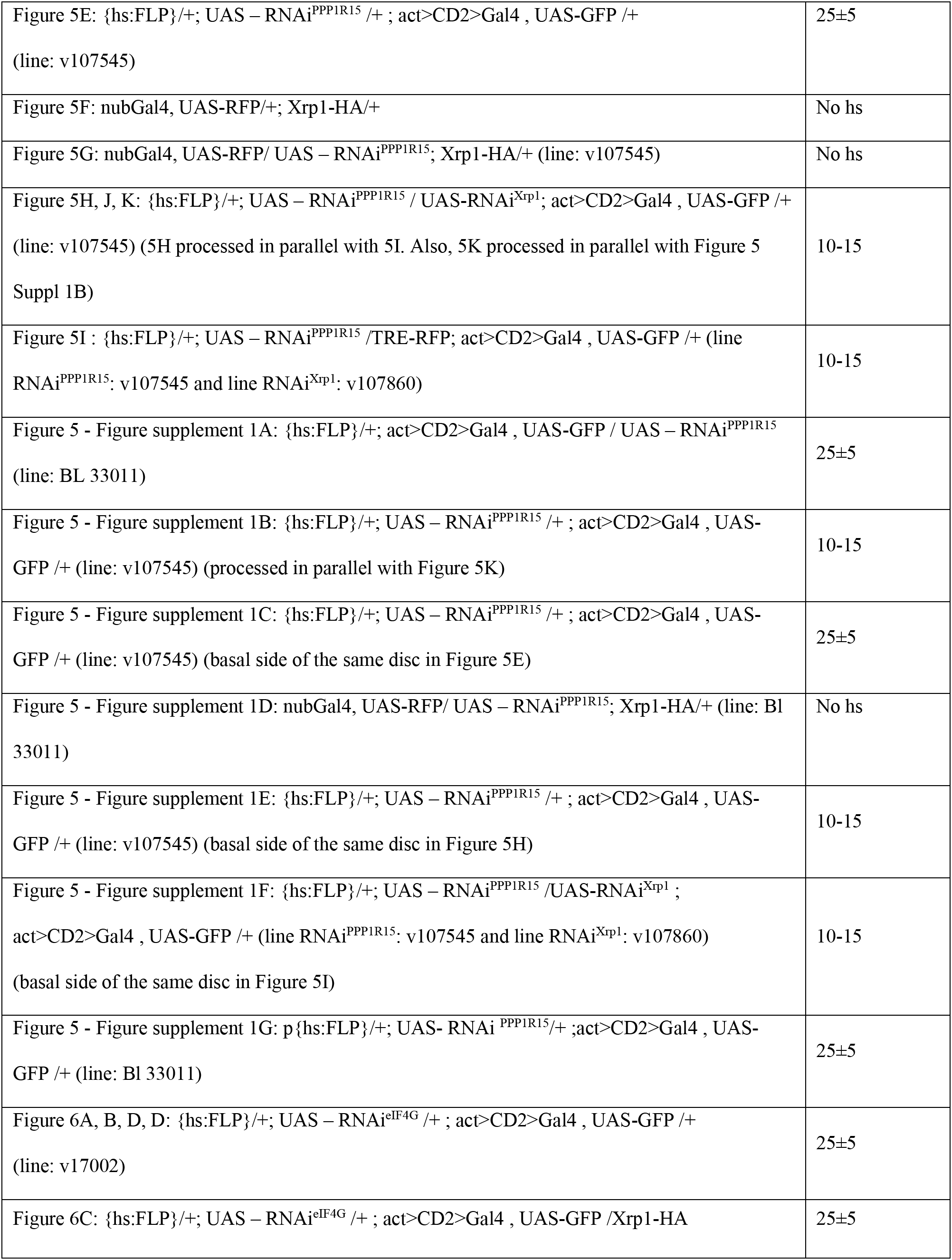
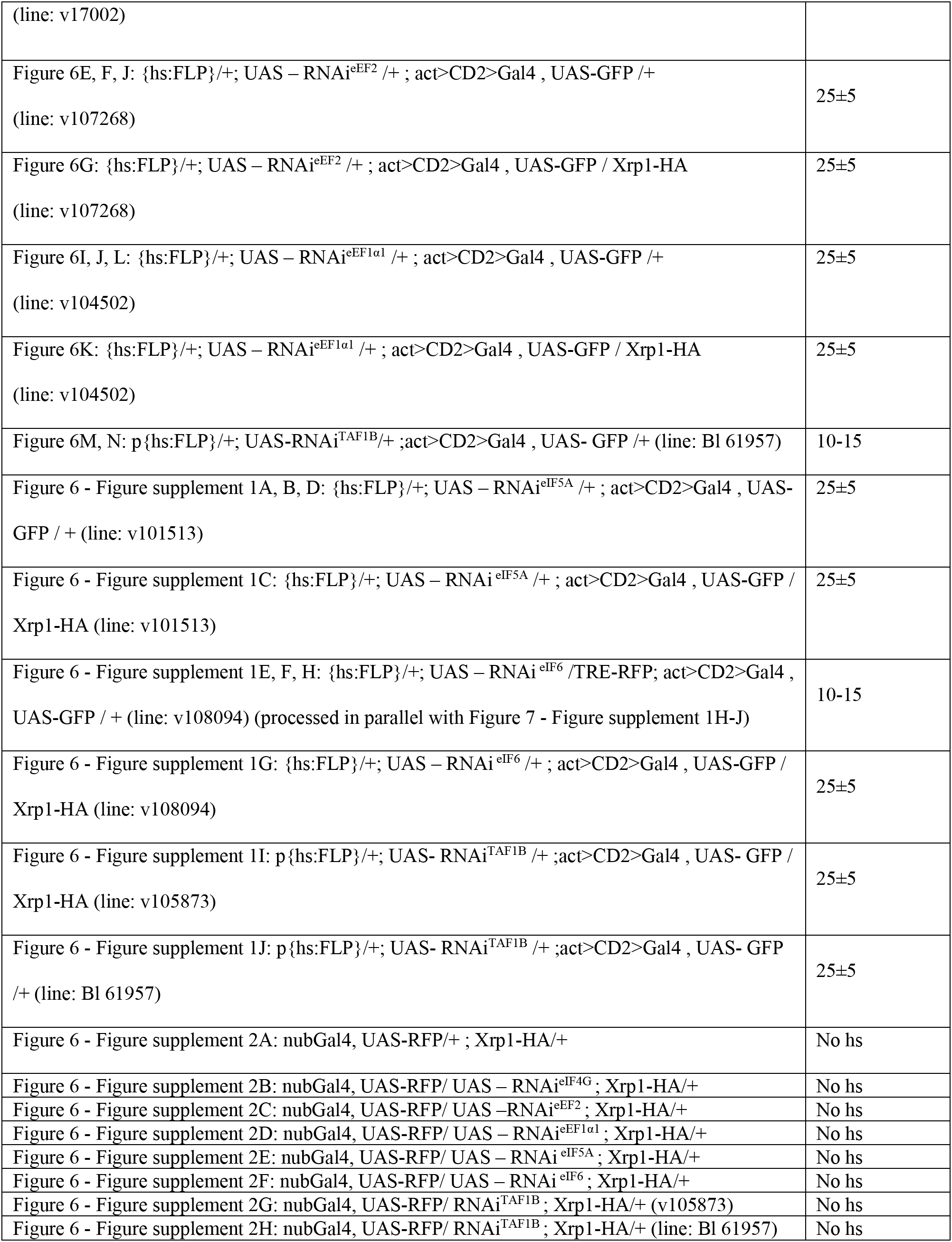
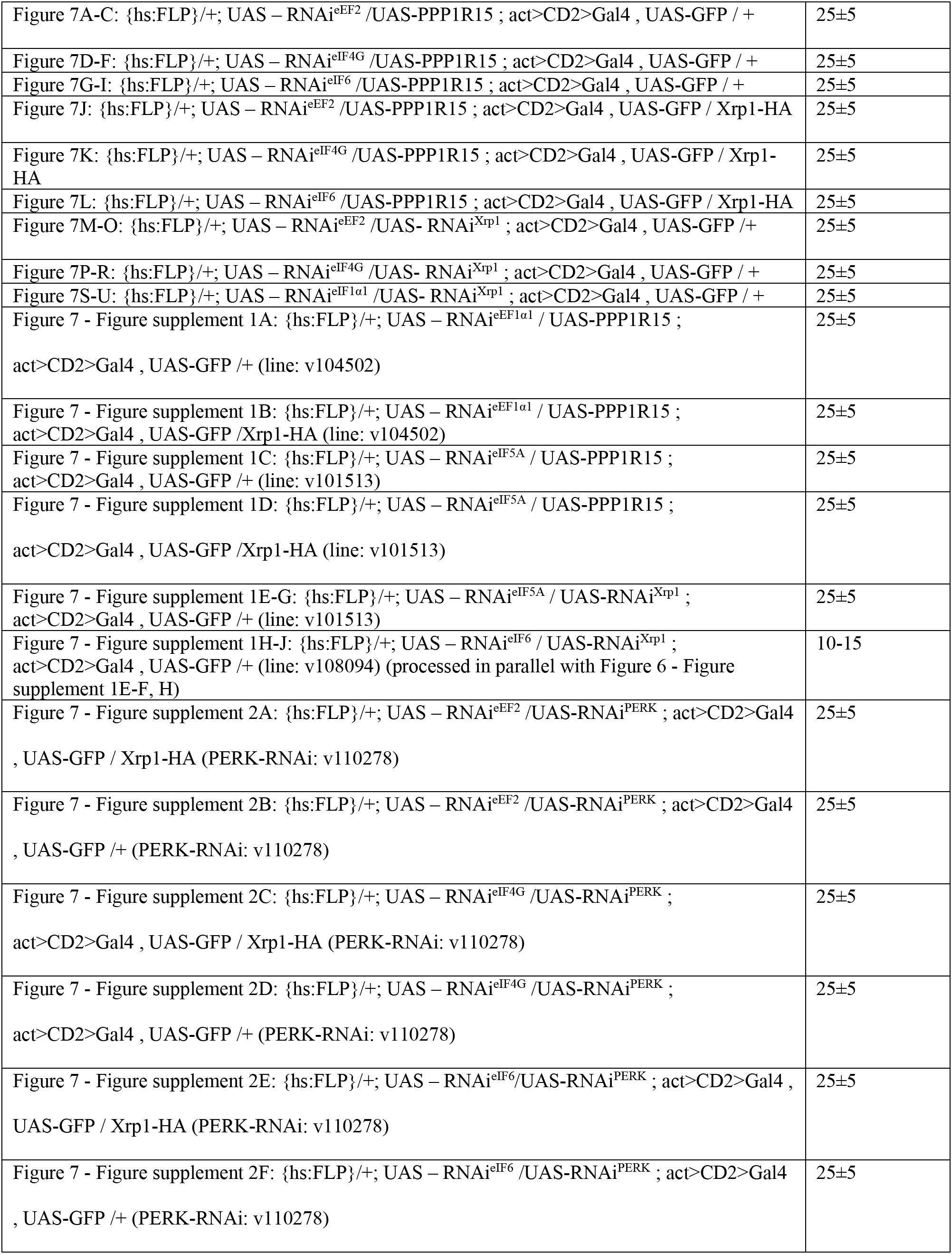
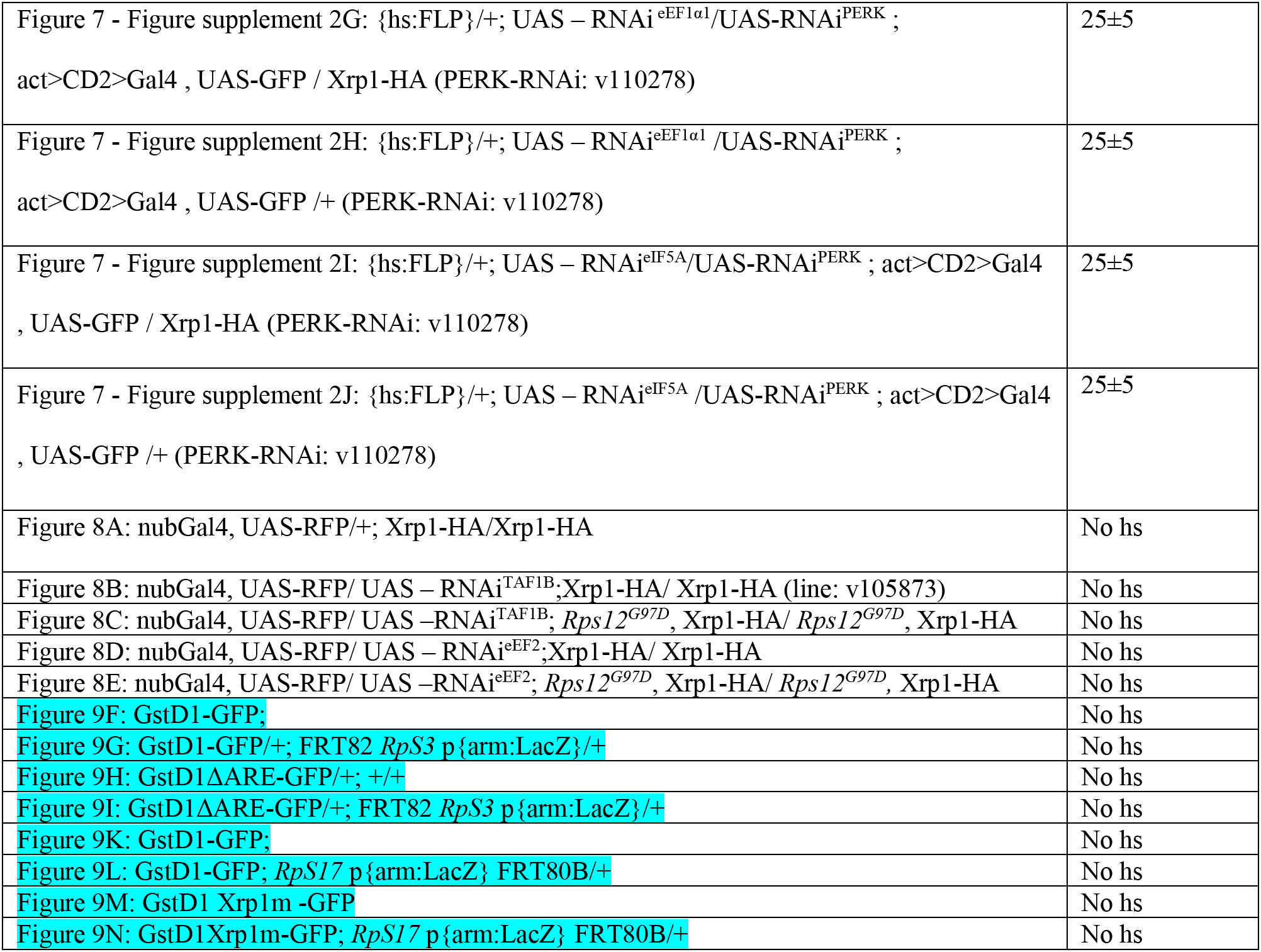
Genotypes and heat shock times of Figures

